# Tissue scale properties of the extracellular matrix regulates nuclear shape, organisation and fate in the embryonic midline sutures

**DOI:** 10.1101/2021.09.03.458867

**Authors:** Diana Alves-Afonso, Allyson Quinn Ryan, Adrian Lahola-Chomiak, Mangal Prakash, Florian Jug, Carl D. Modes, Jacqueline M. Tabler

**Author notes:** These authors contributed equally to this work (Diana Alves-Afonso, Allyson Quinn Ryan).

## Abstract

Balancing self-renewal and differentiation is a key feature of every stem cell niche and one that is tuned by mechanical interactions of cells with their neighbors and surrounding extracellular matrix. The fibrous stem cell niches that develop as sutures between skull bones must balance the complex extracellular environment that emerges to define them with self-renewal and bone production. Here, we address the role for physical stimuli in suture development by probing the relationship between nuclear shape, organization and gene expression in response to a developing collagen network in embryonic midline sutures. This work complements genetic approaches used to study sutures and provides the first quantitative analyses of physical structure in these sutures. By combining multiple imaging modalities with novel shape description, in addition to network analysis methods, we find the early emergence of a complex extracellular collagen network to have an important role in regulating morphogenesis and cell fate. We show that disrupted collagen crosslinking can alter ECM organization of midline sutures as well as stimulate expression of bone differentiation markers. Further, our findings suggest that *in vivo*, skeletal tissues can uncouple the response of the nuclear lamina from collagen mediated tissue stiffening seen *in vitro*. Our findings highlight a crucial relationship between the cellular microenvironment, tissue stiffness and geometry with gene expression in normal development and maintenance of progenitor fate in embryonic sutures.

## Introduction

The balance between stemness and differentiation is an essential feature of organ morphogenesis, where tissue-specific stem cell niches emerge. While biochemical cues are required for specification and maintenance of stem niches, mechanical cues are increasingly found sufficient to control fate acquisition both *in vitro* and *in vivo* where stiffness, composition of the extracellular environment, as well as cellular geometry, modulates potency and fate choice (1). In epithelial niches such as the intestinal crypts, the tight association of polarized cells with a basal matrix confer the physical and geometric stimuli that potentiates stemness 2 3 4 5 6 7. Translating these fundamental insights to mesenchymal tissues and their stem cell niches is challenging, however, as they lack the regular geometric constraints of epithelia and feature complex, dynamic cell shapes and interactions across the tissue and development time. Although *in vitro* mesenchymal stem cells (MSCs) are a powerful model to overcome these challenges and interrogate the cellular role of physical structure and force in fate balance, the predictions made by these studies are rarely probed *in vivo* 8 9 10 11 12. For example, MSCs cultured in substrates of increasing stiffness or geometric confinement increasingly exhibit loss of progenitor identity in favor of skeletal differentiation, suggesting that mesenchymal niches must be protected from tissue stiffening as surrounding cells differentiate, yet few studies have tested this hypothesis directly 8 11 13 14 7 1 15 16.

This proportional relationship between substrate stiffness and differentiation often seen in MSC cultures becomes less clear when considering the suture stem cell (SuSC) niche which form as fibrous, viscoelastic, and flexible joints between the flat bones of the skull 17 18 19 20. These mechanosensitive, connective tissues called sutures give rise to osteoblasts that eventually mediate bone growth in development and repair, while retaining self-renewing progenitors 21 22 23. As might be expected from force-bearing tissues, newborn human sutures are surprisingly stiff with some reports indicating elastic moduli increases from ∼6MPa to ∼400MPa by two years of age 24 25. In contrast, MSCs seeded in 20-40KPa gels exhibit strong upregulation of osteogenic genes such as *Runx2* 8. If tissue stiffening is sufficient to direct skeletal differentiation of stem cells, how can suture stem cells retain their potency as this structure emerges and balances the need for new differentiation?

Just as substrate stiffness influences fate, skeletal differentiation programs are modulated by cell and nuclear geometry as curvature, sphericity, and force propagation across the cell all control MSCs fate balance 26 27 28. The abundant collagen network that propagates compressive and tensile forces thought to stimulate differentiation of new osteoblasts in sutures might do so through geometric constraints imparted by the configuration of this extracellular matrix. However, addressing this hypothesis is challenging without quantitative metrics of SuSC specific gene expression, mesenchymal cell and/or nuclear shapes, alongside matrix organisation during suture formation. Although matrix organization and ossification in response to mechanical loading has been explored at postnatal stages, the developmental relationship between physical stimuli and fate choice has been largely overlooked.

In part, this gap exists because links between transcriptional regulation and mechanoresponsive behaviors relevant to human disease have been unclear, where premature fusion of sutures, known as craniosynostosis, often presents before the first year of life, if not at birth. For example, although cyclic loading has long been found to promote proliferation and differentiation of SuSCs during infancy 29, the most common craniosynostosis patients harbour defects in Fgf, BMP, or Wnt signal transduction 30 31. While the mechanisms through which mechanics might influence these biochemical pathways are not clear 30 31 32, increasing accessibility of patient genome sequencing is determining that previously idiopathic disease arises from mutations in fundamental regulators of cellular mechanics and geometry 33 34 35 36. Of particular interest are a subclass of patients who present with synostosis of midline sutures and harbor mutations in Lamin A (LMNA) 35 37 34, an intermediate filament that modulates tension and stiffness of the nuclear envelope as well as transcription 38 39 40 41 11 42. Although other force transduction networks have been implicated in suture biology 19 33 43, Lamin A orchestrates the nuclear response to Integrin mediated binding of extracellular collagen, an adhesion pathway long known to regulate skeletal differentiation, mesenchymal morphogenesis and, indeed, potency of stem cells in many contexts 44 45 46 47 48 49 50 51 17 52 53.

Intriguing studies emerging from MSC experiments are quickly discovering that LMNA integrates mechanical inputs from integrin mediated adhesion to control gene expression in many ways. Not only important for nuclear shape and genome integrity, LMNA tethers heterochromatin to the nuclear envelope through Lamin Associated Domains (LADs), can bind transcriptional enhancers as well as participate in epigenetic modifications (for example: 54 9 55 42 11 56 57 58 46. Taken together, MSCs respond to stiffer, collagen-rich environments by nucleating contractile actomyosin through Integrin receptor activation which exerts pulling forces on the actomyosin network through cofactors. SUN and Nesprin proteins anchor this contractile network to the nuclear envelope and LMNA coating the interior membrane 41 9 59 60 46 61 47 62 63. Such loading allows LMNA to form stable filaments, leading to stiffening of the nucleus, shape changes and modulation of gene expression, including increased transcription of LMNA itself 11 46 64 26 9 39 41. In contrast, low collagen binding or substrate stiffness releases tension on the nuclear envelope to permit LMNA phosphorylation, releasing it for degradation or binding to transcription factors and enhancer elements 55 9. Again, a discrepancy emerges when considering sutures as they are densely packed with collagen and can far exceed physiological stiffnesses probed in these MSC assays. While laminopathic phenotypes suggest a role for Integrin-Lamin signaling in maintaining suture stemness, how this pathway might regulate fate choice *in vivo* remains confounding.

Here, we explore the role for physical structure, such as collagen and Lamin, in fate choice of prospective suture mesenchyme. To this end, we combine high-resolution microscopy, physical measurements, novel shape description and network analyses to find that suture nuclei are complex and tightly associated where both shape and tissue level organisation can predict identity. We find that the complexity of nuclear shape, spatial organisation as well as skeletal gene expression to be dependent upon Lysyl oxidase-mediated crosslinking of the collagen network that emerges in prospective midline sutures. Surprisingly, such perturbation of physical structure not only influences tissues stiffness but also disrupts LMNA/C accumulation to implicate collagen-lamin signaling in fate balance of skeletal progenitors and suture stem cells. This study reveals a surprisingly early and crucial interplay between the collagen network that defines the suture anatomically and maintenance of progenitors which characterize the suture niche.

## Results

Morphogenesis of the skull vault begins with growth of calvarial bones towards the midline at E13.5, which extend to meet a morphologically distinct mesenchymal condensation at E15.5, that will serve as the suture by the end of fetal development fig1}A) 65. As fusion, whether pathological or not, occurs after this relatively late apposition of frontal and parietal bone fronts when compared to other cranial bone interfaces, midline sutures are most often studied from E15.5. At these stages, canonical markers used in developmental studies, such as Twist, Msx2, Gli1 or Axin2 are not exclusively expressed in sutures as they can also be found in adjacent cell populations of bone, dermis or meninges 66 67 68 69. Similarly, both stem cell markers and skeletal progenitors that we found highly represented in bulk RNA sequencing of E13.5, 14.5 and E15.5 sutures marked both suture and bone Figure 1 A, Figure supplement I. Finding no suitable suture marker *per se*, we instead sought to understand when sutures begin specialising and compared the morphologically distinct E15.5 sutures with E14.5 and 13.5. Principle Component Analysis (PCA) revealed significant changes in gene expression between E13.5 and 14.5, as well as between E14.5 and E15.5, suggesting dramatic shifts in gene expression throughout the time points explored and earlier than is often studied in suture development Figure 1 C. Differences between datasets were further determined by differential gene expression (DGEs) and Gene Ontology (GO) analyses and revealed that E15.5 sutures are considerably distinct from E13.5 prospective sutures, where genes involved in physical structure, such as ECM composition and assembly, are increasingly enriched in E14.5 and 15.5 sutures (compare Figure 1 D Figure supplement II and Figure supplement III). These data suggest that by E14.5, prospective sutures have initiated construction of its fibrous ECM network and must begin responding to a new mechanical environment early in skull development.

**Figure 1.**
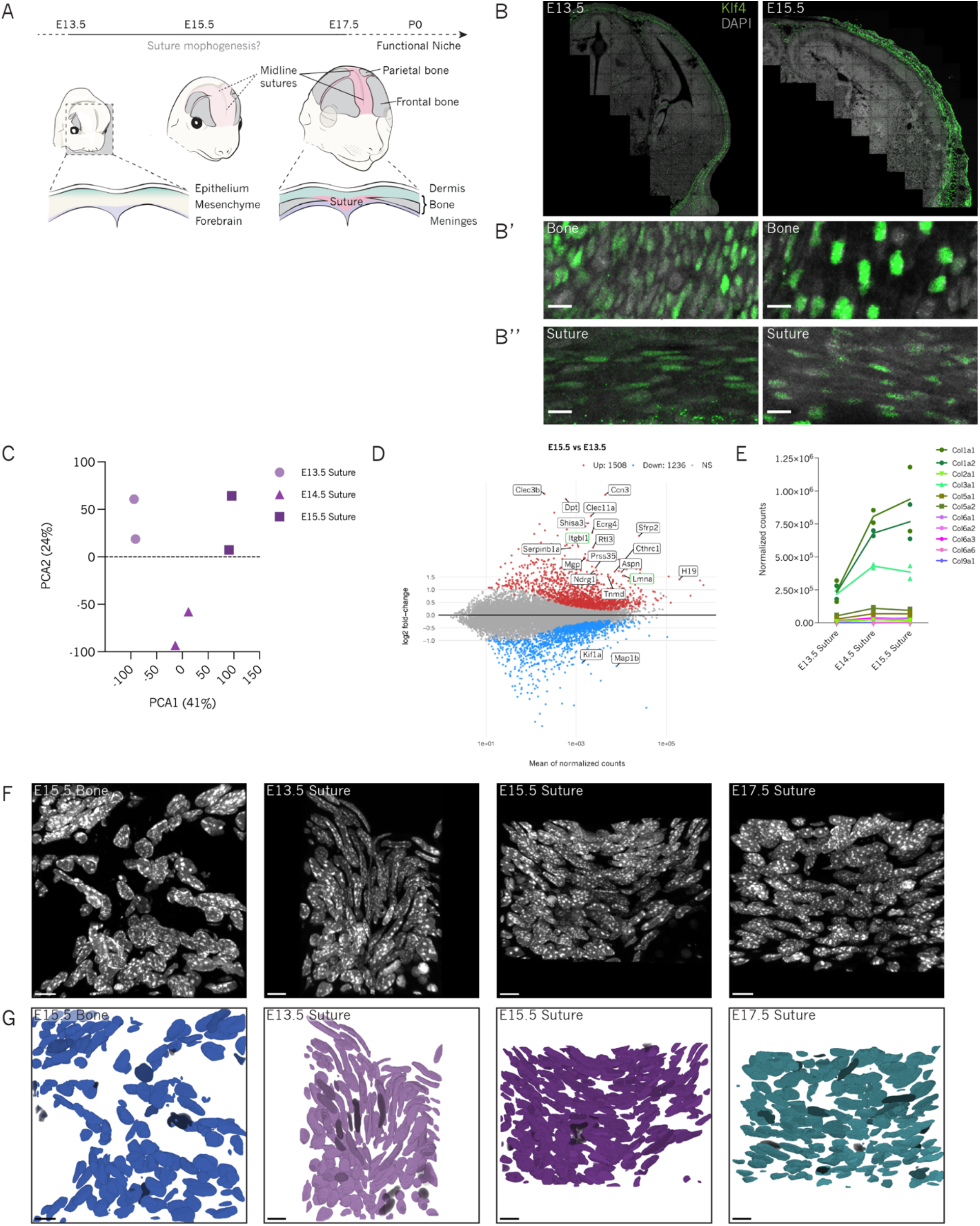
Genetic approaches are not sufficient to specifically label the embryonic midline suture. A) Schematic showing the development of the skull vault including frontal and parietal bones and cranial sutures. B) Coronal section of E13.5 and E15.5 heads shows KLF4 positive cells in green. Scale bar 100 µm. B’) Enlarged view of frontal bone region shows cells positive for KLF4 at both developmental stages. Scale bar 10 µm. B’’) Enlarged view of midline suture at E13.5 and E15.5 shows KLF4 positive cells in the suture. Scale bar 10 µm. C) PCA analysis of E13.5, E14.5 and E15.5 midline sutures. Samples were projected onto the 2D plane such that they spread out in the two directions that explain most of the differences. The x-axis is the direction that separates the data points the most and the y-axis is the direction that separates the data the second most. Biological replicates in each dataset are indicated in the same shape and color. D) The log2 fold change for E15.5 vs. E13.5 comparisons is plotted on the y-axis and the average of the counts normalized by log2 is shown on the x-axis. Each gene is represented with a dot. Upregulated genes are shown in red and downregulated genes are shown in blue. Top 20 the of differentially expressed genes per condition have been labeled. E) Expression of collagen encoding genes by midline sutures at E13.5, E14.5 and E15.5. Fibrillar collagens are shown in different shades of green and fibril-associated collagens are shown in different shades of pink. F) 3D rendering of confocal z-stacks of frontal bone and midline sutures at different developmental stages. Tissue sections were stained with DAPI shown in gray. Scale bar 10 µm. G) 3D renderings of segmented frontal bone and midline suture nuclei. Scale bar 10 µm.

The most highly represented ECM protein expressed in E14.5 and E15.5 midline sutures were fibrillar collagens including *Cola1a1, Col1a2* and *Col3a1*, which will assemble to form Type I, Type II, Type III fibrillar collagen Figure 1 E. As such, collagens which not only mediate tissue stiffness, but also govern cell and nuclear geometries, increased across developmental time, we asked whether morphological analyses could be used to infer cell fate decisions while simultaneously providing a geometric characterisation of the suture niche as it is established. Cell and nuclear morphology have long been used as indicators of cell state as changes in morphology are linked to alterations in cell fate or function. For example, different cell geometries modify actomyosin contractility, which directs MSCs to differentiate into osteoblasts or adipocytes *in vitro* 14. Unlike polarised epithelial niches, where the tight association of polarised cells allows individual cells to be tracked through membrane or junction labels, suture mesenchyme lacks these distinctions and individual cell morphologies can only be parsed by mosaic cell labelling Figure supplement IV. Such sparse membrane labelling experiments will be important for probing the cell biology of individual cells but understanding tissue level control of cell fate requires whole tissue analyses. As nuclear shapes are far simpler to distinguish, we chose to analyse and map the geometry of nuclei at early middle and late stages of suture development from volumetric stacks of DAPI stained coronal sections Figure 1 F Figure supplement V A. DAPI labelled nuclei were also very complex, including some which were very tightly associated, featuring striking convexities and concavities. The complex nuclear shapes and packing we found throughout early development of these sutures presented a significant challenge in quantitative approaches as existing image segmentation tools performed poorly where manual corrections required significant time investment.

### Nuclear geometry is sufficient to isolate populations in the cranial mesenchyme

To improve accuracy of image segmentation we appealed to Deep Learning Methods and developed a novel algorithm for generating semi-automated segmentation masks from our suture cross-sections Figure 1 G Figure supplement V B and compared these with nuclei from differentiated E15.5 frontal bones as a control reference. These volumetric nuclear segmentations then provided a means for quantitatively describing nuclear geometry through shape description targeted at particular features, such as sphericity, which is classically determined as a ratio of object volume and surface area 70 and is typically calculated using polygonal meshes as proxies for these voxelated-object. However, well-known mesh generating methods (e.g. convex hull or marching cubes) frequently lead to volume overestimation and surface area underestimation due to the requirements of existing algorithms Figure 2 A. It is possible to balance these errors using complex closed curve fitting methods (e.g. β-spline fitting) but, unfortunately, such approaches are computationally expensive and therefore not fit for high-throughput studies. Given the apparently complex nuclear surfaces of suture cells, we derived an alternative definition of sphericity (see Methods) that is purely dependent on object volume rather than surface estimations allowing us to circumvent previous methods. We found that nuclear sphericity values were significantly different in tightly-associated suture mesenchyme when compared to meningeal cells and could be distinguished computationally Figure supplement VI. Additionally, we note that the mean and standard deviation of suture nuclear sphericity decreases over developmental stages Figure 2 C, D, which implies that the shape of suture nuclei is remodeled over the course of development to generate flatter nuclei when compared with differentiated control osteoblasts of E15.5 frontal bones.

**Figure 2.**
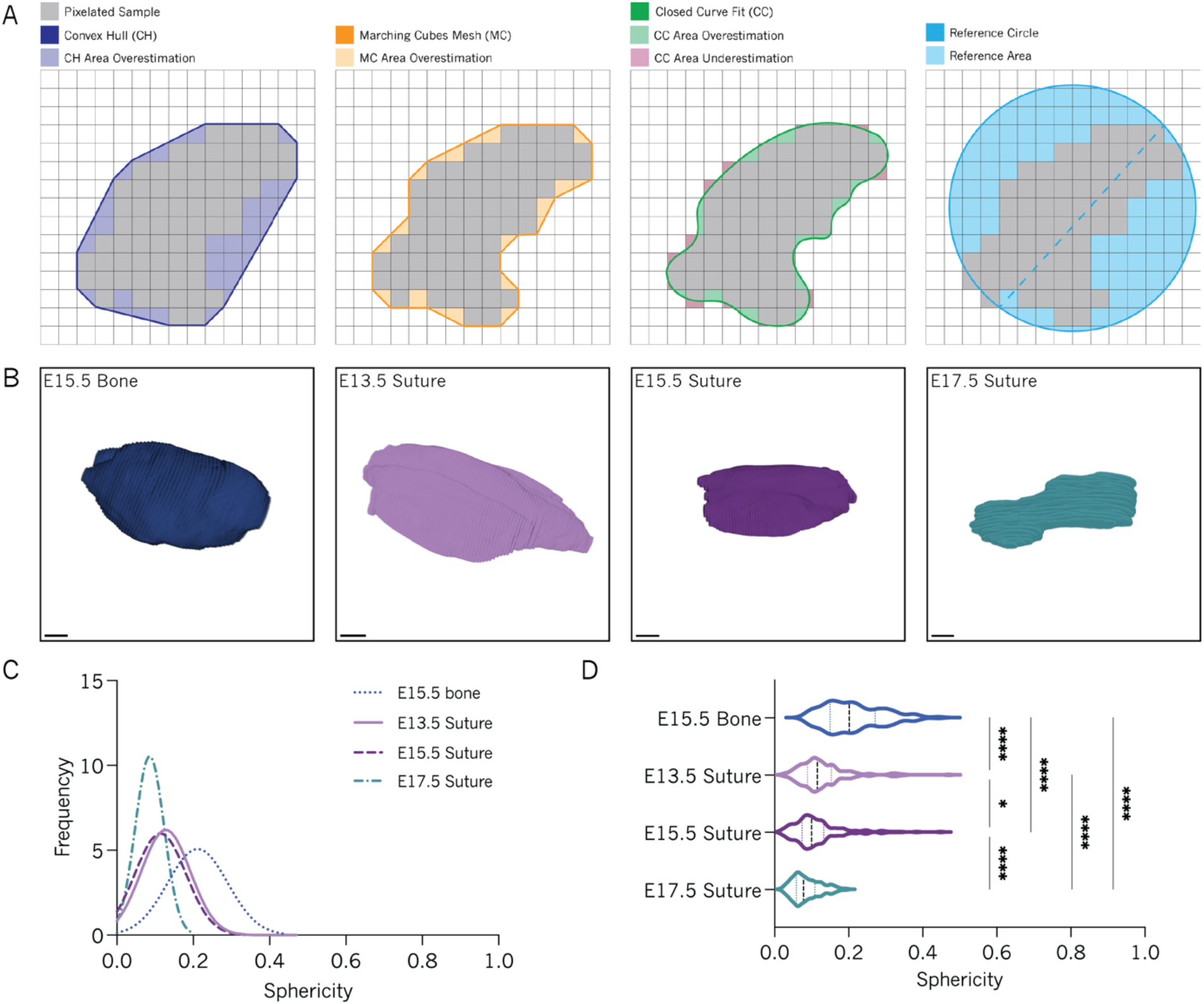
Suture nuclei become progressively flatter in comparison to adjacent mesenchymal cell populations. A) Schematic representing common surface area estimation algorithms (Convex Hull, left; Marching Cube mesh generation, center left; closed curve spline fitting, center right) and the alternative reference method proposed here (right). B) 3D rendering of a randomly selected nucleus from each one of the analyzed datasets. Scale bar 10 µm. C) Frequency distribution of surface area independent sphericity measurements for the different tissues and timepoints. D) Violin plot of volume-based sphericity measurements for segmented E15.5 frontal bone nuclei (N=2, n=94) and E13.5 (N=2, n=180), E15.5 (N=2, n=196) and E17.5 (N=2, n=213) midline suture nuclei. N represents the number of animals, n represents the number of individual nuclei. Kruskal-Wallis test was applied to estimate p-value. *p<0.05, ****p<0.0001.

Though assessing nuclear sphericity is sufficient to computationally distinguish presumptive meningeal nuclei from suture nuclei at early developmental stages, sphericity is a poor descriptor when so many surface irregularities can be observed across nuclei. To access more of the shape information (i.e. geometrically distinct properties) of suture nuclei, we chose to extract topological skeletons of individual nuclei 71. This again allowed us to avoid the need to work with cumbersome, explicit 2D surface representations in 3D space, by generating a reduced representation of the surface variability of 1D voxelated curves in 3D space Figure 3 A, B. We used the standard *N*_2_6 voxel neighborhood to parse topological skeletons into a series of branches, which allows us to use each branch as a proxy for a surface irregularity (e.g. concavity). We observed a higher degree of variability in both skeleton branch count and length in suture nuclei when compared to osteoblast nuclei suggesting that SuSCs form with greater surface complexity when compared to differentiated cells of reference bone. Although branch number increased as sutures develop, branch length decreased with developmental time Figure 3 C, D, together indicating that nuclear features, such as surface concavities, may be actively regulated and dynamic.

**Figure 3.**
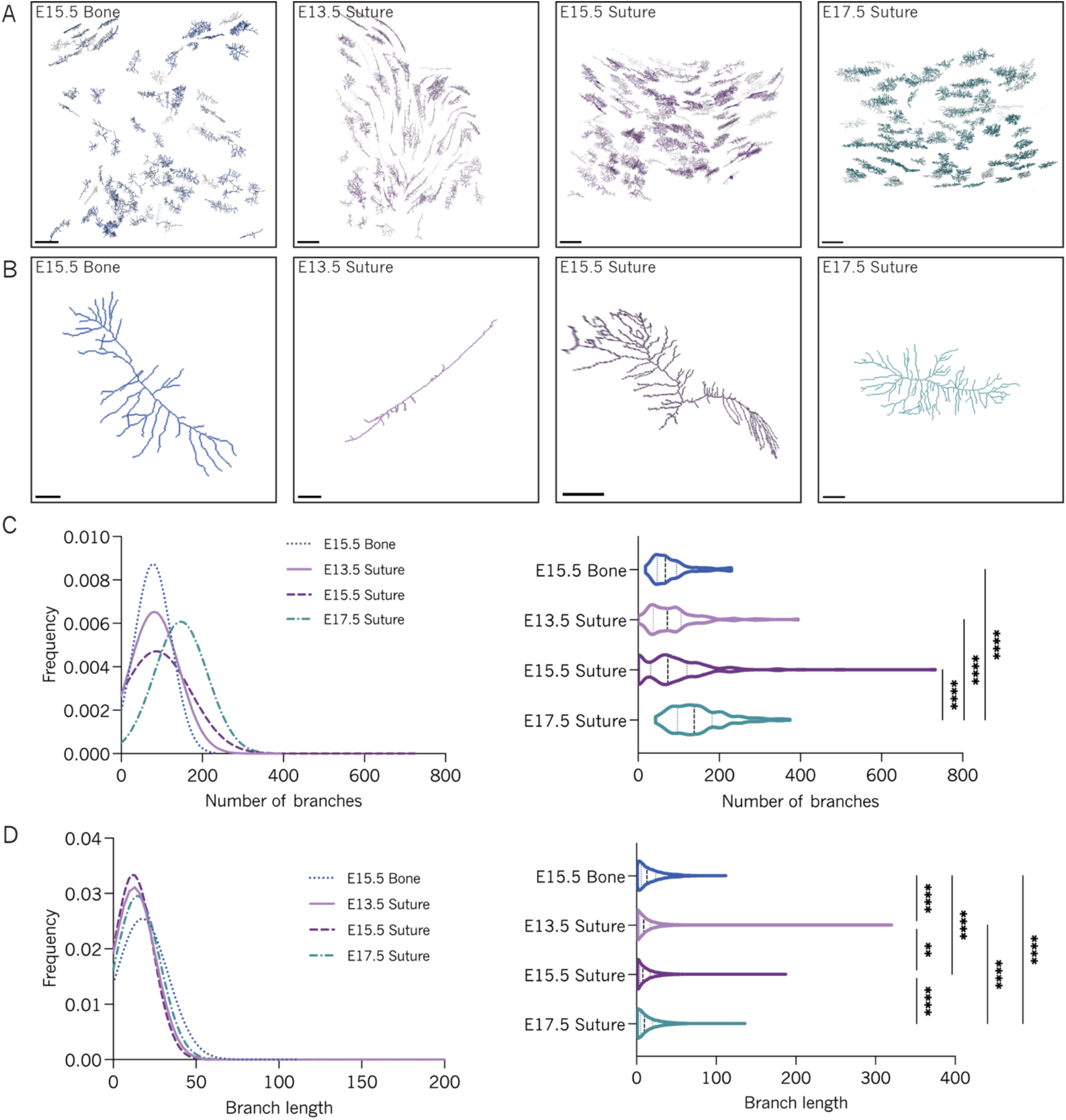
Topological skeletons are distinct between stages of suture development as well as from those of bone. A) 3D rendering of topological skeletons of E15.5 bone and E13.5, E15.5 and E17.5 suture. Scale bar 200 voxels. B) 3D rendering of a randomly selected single topological skeleton for each of the studied timepoints. Scale bars fro left to right 30, 40, 30, 40 voxels. C) Frequency distribution (left) and violin plot (right) of topological skeleton branch number for E15.5 frontal bone nuclei (N = 2, n = 94) and E13.5 (N = 2, n = 180), E15.5 (N = 2, n = 196) and E17.5 (N = 2, n = 213) midline suture nuclei. D) Frequency distribution (left) and violin plot(right) of topological skeleton branch length (see C for N, n values).

To further understand the unique tissue level spatial aspects of the developing midline suture, we generated networks representing nuclear neighbor relationships. Such networks are usually created using an object’s center of mass for the network node proxy as the objects of interest tend to have regular shapes. However, as is clear from single-cell level shape characterization, suture nuclei are geometrically complex and frequently feature highly irregular shapes. Therefore, to incorporate sensitivity to such shape complexity into a tissue network, we instead used the explicit nuclear shapes calculated from our image segmentations. Network edges which are defined as the connections between two nuclei and represent neighbor associations, were accordingly found by identifying the minimal distance between nuclear surface boundaries. To examine how networks differed across both tissues and stages, we generated networks over a range of distance thresholds where two edges fall below a given distance threshold, both objects - or nuclei are included in a neighborhood network Figure 4. At very short distance thresholds presumptive suture, meningeal and dermal nuclei segregate separately in our network analyses although this segregation is lost at distances greater that 5-10 µm depending on developmental stage Figure 4 A-E. Such segregation of clusters based on surface distances suggests that each tissue-specific cell type has distinct organisation. The giant component fraction, which is defined as the largest connected cluster of nuclei within a network, will equal 1 when each nucleus within the image is connected and can be used to quantify how tightly associated objects are within an ROI. We found the giant component fraction to reach saturation at shorter distance thresholds in sutures when compared to reference bone nuclei, indicating tighter association of cells across prospective suture mesenchyme. However, the giant component fraction reached saturation at increasingly greater distance thresholds over developmental time, where E13.5 and E15.5 suture networks reached saturation at 5 µm but E17.5 suture and E15.5 bone saturation require approximately 10 µm and 15 µm respectively Figure 4 E. The same ordering of metric saturation (i.e. E13.5 suture reaches saturation first, followed by E15.5 suture, then E17.5 suture and finally E15.5 bone) seen in the giant component fraction can be observed in the mean of the networks’ clustering coefficients Figure 4 E, F. The standard deviation of clustering coefficients were in the inverted order where cluster incorporation occurred over shorter distances and with decreased variability suggesting tighter and more regular association of cells across the entire suture mesenchyme when compared to bone Figure 4 G. Another metric of connectivity that is indicative of mutual neighbor density can be calculated from the number of closed paths across neighbor objects within a network, represented here as number of cycles. We observed greater cycle numbers at shorter distance thresholds in early sutures at E13.5 and E15.5, when compared to E17.5 suture mesenchyme or bone Figure 4 H. These data suggest that suture midline mesenchyme undergoes reorganization of neighborhood spacing of nuclei as the suture matures.

**Figure 4.**
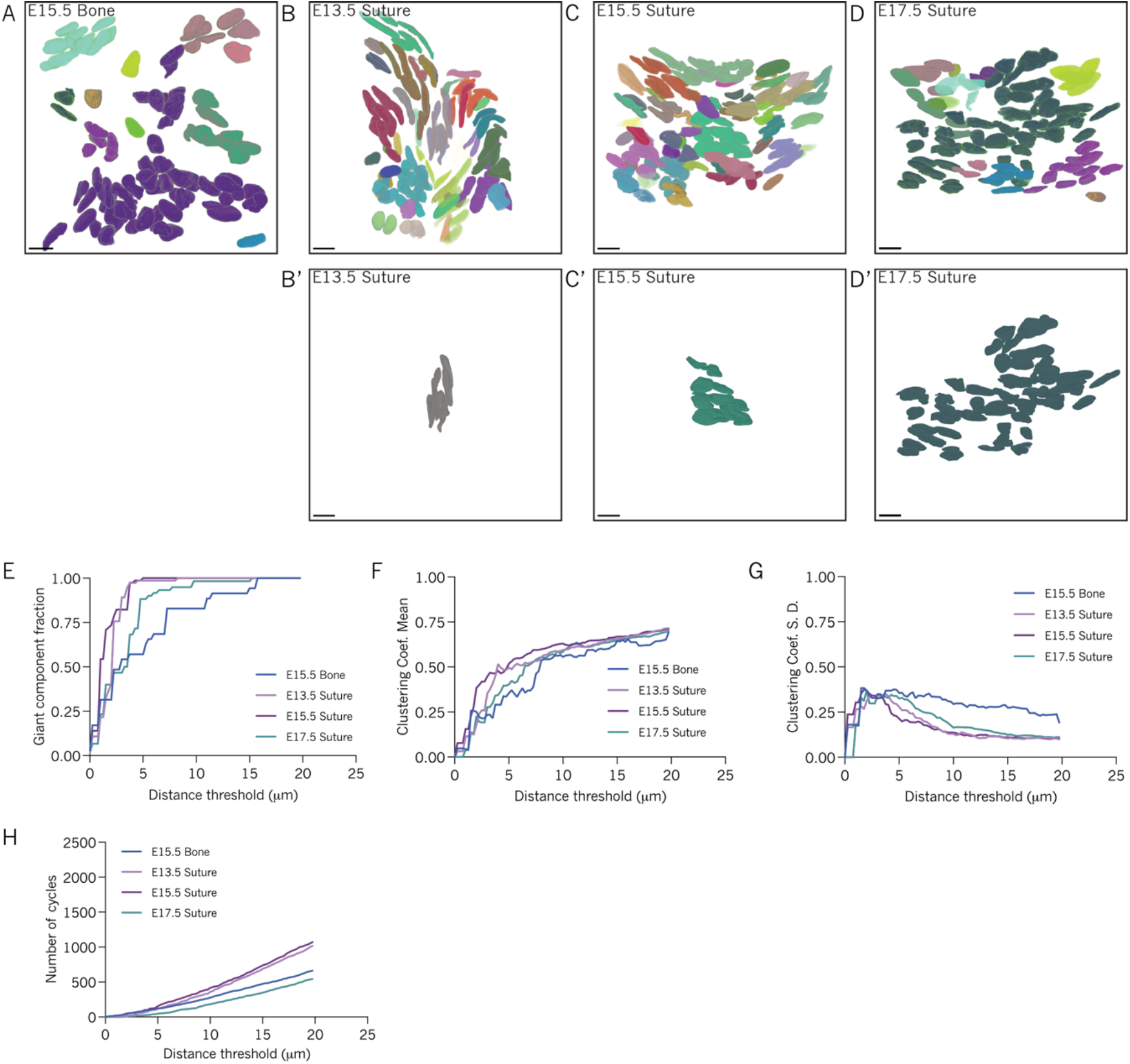
Suture nuclear neighbor relationships and network metric progressions indicate early suture nuclei to be tightly associated. 3D renderings of nuclear clusters for A) E15.5 frontal bone and B) E13.5, C) E15.5 and D) E17.5 midline suture nuclei. 3D renderings of an isolated nuclear cluster for B’) E13.5, C’) E15.5 and D’) E17.5 corresponding to potential suture nuclei. E) Line plot of giant component fraction for E15.5 frontal bone and E13.5, E15.5 and E17.5 midline suture minimal distance-based (MDB) nuclear networks. Line plots of F) clustering coefficients mean and G) standard deviations for E15.5 frontal bone and E13.5, E15.5 and E17.5 midline suture MDB nuclear networks. H) Line plot for cycle basis counts for E15.5 frontal bone and E13.5, E15.5 and E17.5 midline suture MDB nuclear networks.

### Fibrillar collagens are assembled in early suture development

As nuclear shapes can be regulated by collagen through Integrin-LINC pathway 11, we wondered whether the collagens we found to be highly represented in these sutures Figure 1 E were also assembling into networks that may explain nuclear shape and organisation in these sutures. Therefore, we assessed collagen fibrillogenesis in E15.5 sutures, a stage which marked the most significant increase in collagen expression, via transmission electron microscopy (TEM). This revealed a densely packed network of extracellular collagen fibrils aligned in both anterior-posterior (AP) and medio-lateral (ML) axes of the skull Figure 5 A, A’. AP oriented fibrils appear as circles in coronal sections while ML oriented fibers are seen in the longitudinal plane. Remarkably, signs of ECM maturation and collagen fiber formation are observable from early stages of suture development at E13.5 Figure 5 B. These data suggest that a complex extracellular environment is established in the earliest stages of skull morphogenesis, much earlier than has been previously described 19, 20. While the collagen network observed in electron micrographs appeared extensive, we sought a quantitative description of collagen structure in developing sutures to better associate the cellular organisation of the suture with fate choice. As tissue level characterizations using TEM is technically challenging because it is optimized for nano-scale imaging, we turned to polarized light microscopy (PLM), which uses the optical properties of materials/tissues, for label-free detection and calculation of tissue scale organization 72, 73. More specifically, PLM takes advantage of birefringence, an optical property that collagen possesses, where anisotropic refractive indices are used to extract measurements for collagen orientation in whole tissues. PLM images from E13.5, 15.5 and 17.5 sutures were processed such that collagen orientation is represented by distinct hues Figure 5 C. To determine the predominant orientation angle(s) within the suture, PLM images were filtered for areas of collagen fibril enrichment and alignment to remove background signal arising outside of the tissue (Figure 5 C; see Methods). We then selected suture ROIs within images from which angle measurements were calculated Figure 5 C, D. The angle quantification revealed a progressive alignment of collagen fibrils within the suture region as the tissue develops Figure 5 D. At earlier stages there did not appear to be a preferred orientation, while in later stages (E15.5 and E17.5) fibers were preferentially aligned along the mediolateral axis Figure 5. These data show geometric changes to the collagen network that may underlie nuclear organisation, and that later enforce distinct neighbor associations of suture cells relative to those in dermis or meninges.

**Figure 5.**
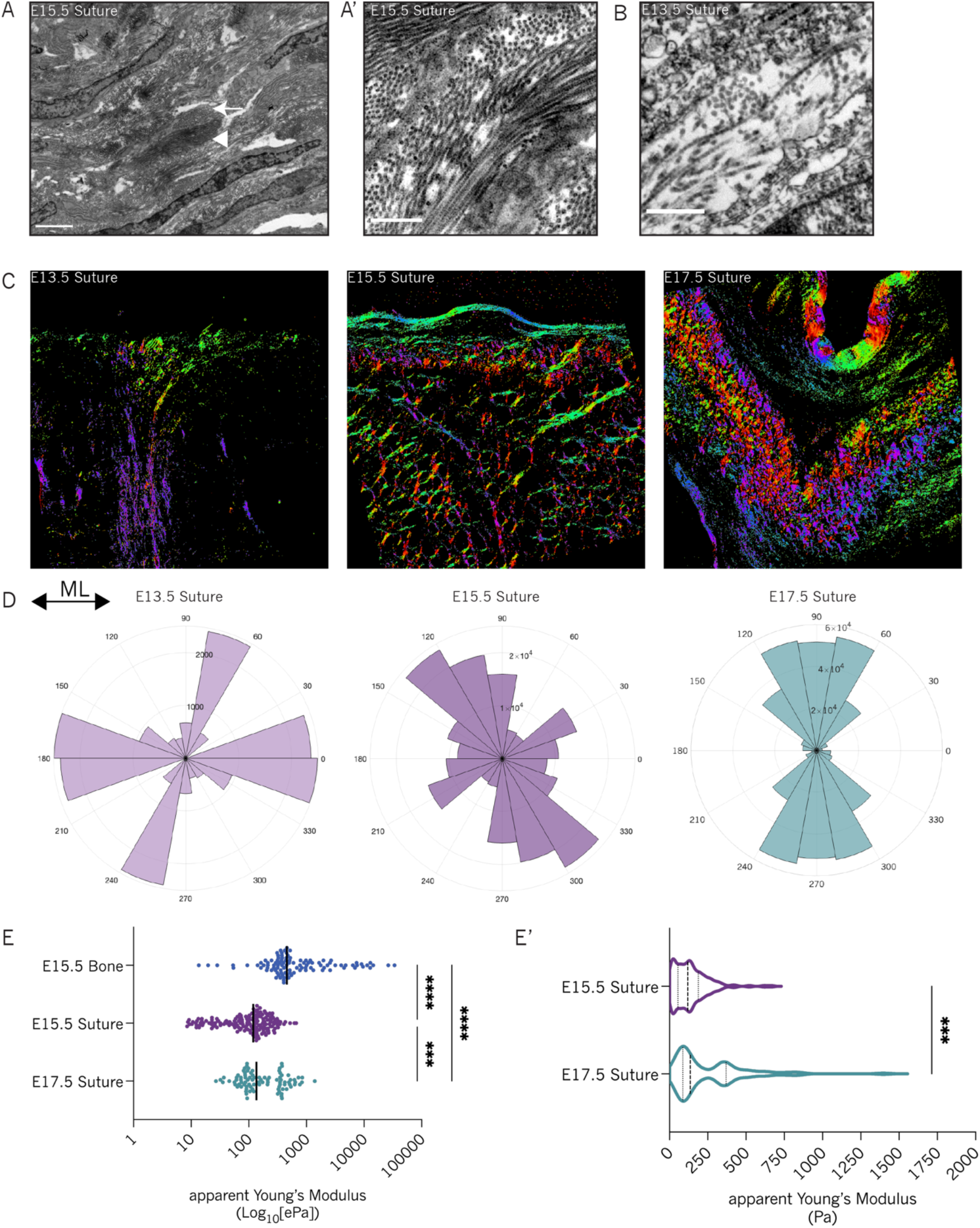
Embryonic midline sutures are rich in fibrillar collagen and are orders of magnitude softer than frontal bone. A) TEM view of E15.5 suture shows densely packed collagen fibrils in the extracellular space. White arrowhead indicates collagen fibrils oriented along the longitudinal plane. White arrow points to collagen fibrils oriented along the anterior-posterior axis. Scale bar 0.5 µm. A’) Enlarged view of E15.5 suture under TEM shows collagen fibrils with different orientations. Scale bar 0.5 µm. B) Enlarged view of E13.5 suture under TEM shows some bundles of collagen fibers. Scale bar 0.5 µm. C) Representative polarised light microscopy images for E13.5, E15.5 and E17.5 midline sutures (cyclic LUT represents collagen fibril orientation). ML, medio-lateral. D) Rose plots showing the distribution of collagen orientation angles in E13.5, E15.5 and E17.5 suture regions. E) Scatter plot shows tissue bulk stiffness for E15.5 frontal bone (N=8, n=105), E15.5 (N=11, n=166) and E17.5 suture (N=5, n=90). N represents the number of animals, n represents the number of individual force curves. Black line indicates the median. Kruskal-Wallis test was applied to estimate p-value. ***p<0.0002, ****p<0.0001. E’) Violin plot shows that the distribution of calculated apparent Young’s Modulus is statistically different between E15.5 and E17.5 midline suture.

The fibrillar collagens we found to be expressed in developing midline sutures are the primary extracellular matrix structures that resist shear strain 74. A recent study has reported that differences in tissue specific stiffnesses are correlated with abundance of collagen and, therefore, we measured tissue bulk stiffness in the embryonic midline suture. For this, we developed an Atomic Force Microscopy (AFM) protocol that allowed us to measure the apparent Young’s Modulus of tissues in the suture at E15.5 and E17.5 and compared these to values recorded in E15.5 bone, structures which mineralise to become significantly stiffer than sutures. As expected, E15.5 bone was found to be far stiffer than midline sutures at both E15.5 and E17.5 where the apparent Young’s Modulus of E15.5 bone measures at 2201Pa (± 479.2 SEM), at least an order of magnitude greater than that measured in developing sutures, which measure 135.6Pa(± 8.674 SEM) and 251.6 Pa (± 25.75 SEM) at E15.5. and E17.5, respectively Figure 5 D. This significant difference in bulk tissue stiffness values between suture (undifferentiated) and bone (differentiated) is consistent with previous observations suggesting that skeletal differentiation is promoted by high tissue stiffness 8 41 9 64. Nevertheless, we observed a 1.8 fold increase in the apparent Young’s Modulus of the suture from E15.5 to E17.5 Figure 5 D’.

### Role of collagen crosslinking in midline suture development

If the emergent collagen network is responsible for cell fate, we predicted that disrupting the mechanical properties of this network would cause premature differentiation. To address this hypothesis we considered the additional structural proteins that lock collagen fibrils together through modifications such as crosslinking. For example, extracellular collagen fibers that form from exported collagen fibrils are often strengthened by covalent cross-links promoted by Lysyl oxidase (LOX). While LOX is necessary for canalising the collagen network of bone that will later mineralise, the role of LOX in sutures has never been addressed, although, our bulk RNA sequencing reveals that LOX is, indeed, expressed in the suture Figure supplement VII C. To ask whether the mechanical properties of collagen are mediated by LOX crosslinking, we chose to inhibit the activity of LOX by feeding pregnant dams with β-Aminopropionitrile (BAPN), an irreversable inhibitor of LOX activity 75 76 77 78 79 80 81. To avoid disrupting early developmental requirements for LOX, we began feeding dams 11.5 days post conception at a time after the mesenchyme that will give rise to the skull is resident over the cortex, and flanking calvarial bones are being specified Figure supplement VII A. Inhibition of LOX activity through BAPN has previously been reported to impact collagen fiber formation and shape leading to altered material properties across tissues 79 and long term defects in endochondral bone 78 80 81. Therefore, we first asked whether BAPN treatment induced defects in collagen fibrillogenesis. To this end, we prepared BAPN samples for TEM and screened the midline suture to quantitatively compare collagen fiber morphology in the WT with BAPN treated midline sutures. We measured both the cross-sectional area, circularity and aspect ratio of fibrils using FiJi. Area measurements confirm the that the average size of collagen fibrils in BAPN embryos (E15.5 Mean = 1947 nm^2^ ± 30.52 SEM; E17.5 Mean = 2011 nm^2^ ± 28.09 SEM) is significantly higher than WT fibers (E15.5 Mean = 1603 nm^2^ ± 16.00 SEM; E17.5 Mean = 1631 nm^2^ ± 17.17 SEM) Figure 6 B. We also found significant changes in fibril circularity and aspect ratio in E15.5 BAPN treated embryos Figure 6 B, Figure supplement VII B, confirming that BAPN mediated LOX knockdown confers fibrillogenesis defects *in vivo*. Importantly, the distribution of area values in BAPN treated sutures when compared to WT, highlights the heterogeneity of collagen fibril morphology found in the BAPN treated embryos.

**Figure 6.**
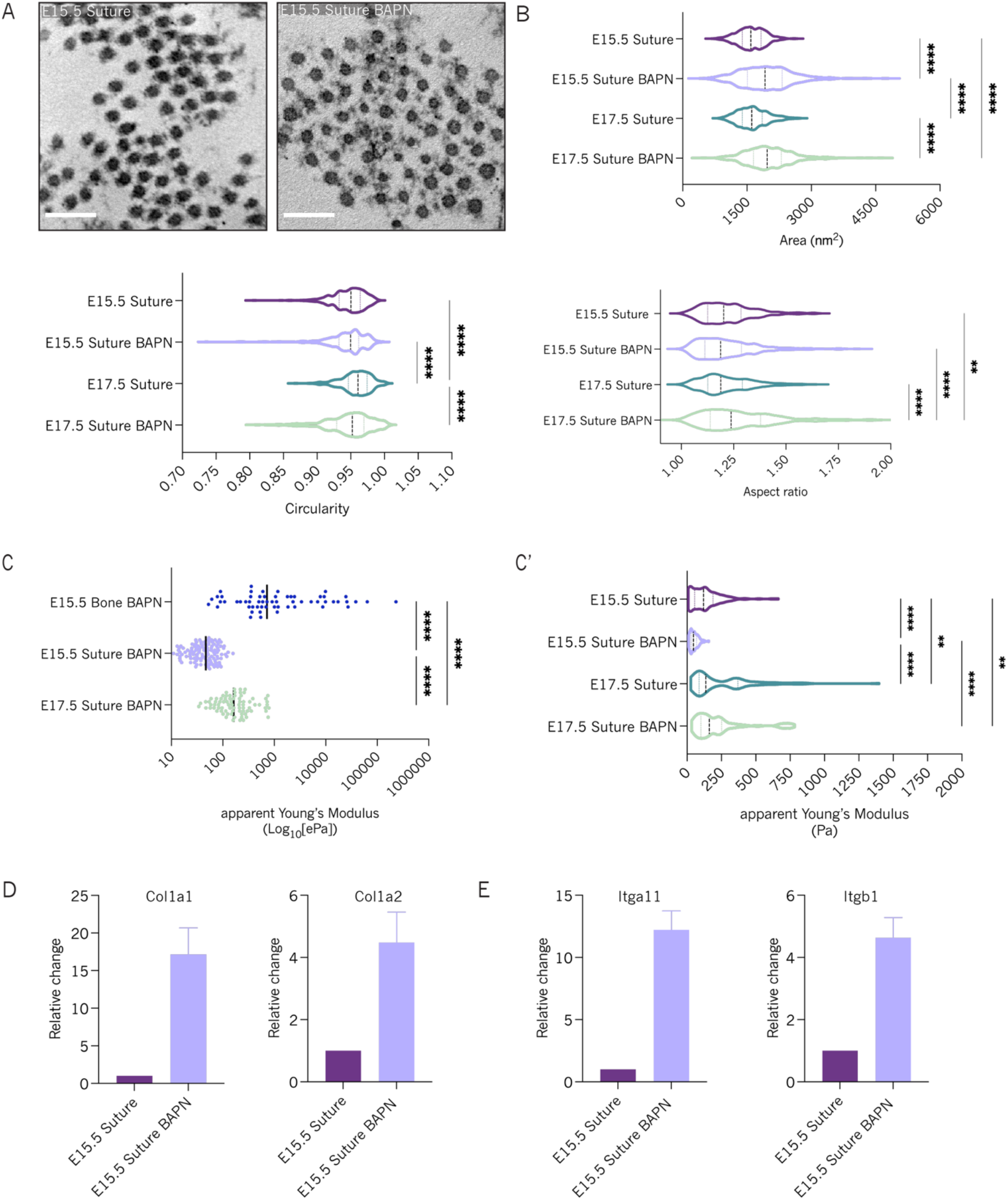
Inhibition of collagen crosslink decreases tissue bulk stiffness. A) TEM view of collagen fibers in the suture of WT and BAPN treated E15.5 embryos. Scale bar 200nm. B) Violin plots comparing area (nm^2^), aspect ratio and circularity of collagen fibers in E15.5 Suture (N=2, n=400), E15.5 Suture BAPN (N=2, n= 409), E17.5 Suture (N=1, n=402) and E17.5 Suture BAPN (N=2, n=385) embryos. N represents the number of animals, n represents the number of analyzed collagen fibers. Dashed line indicates the median. Kruskal-Wallis test was applied to estimate p-value. *p<0.05, ****p<0.0001. C) Scatter plot shows tissue stiffness for E15.5 Bone BAPN (N=5, n=59), E15.5 Suture BAPN (N=9, n= 117) and E17.5 Suture BAPN (N=6, n=79). N represents the number of animals, n represents the number of individual force curves. Black line indicates the median. Kruskal-Wallis test was applied to estimate p-value. ****p<0.0001. C’) Violin plot shows the distribution of calculated apparent Young’s Modulus values for E15.5 and E17.5 WT and BAPN midline suture. Kruskal-Wallis test was applied to estimate p-value. ****p<0.0001. D) Expression levels of *Col1a1* and *Col1a2* genes, which encode for Type I collagen, are increased in the midline suture of BAPN treated animals relative to WT. E) qPCR measurements show increased levels of expression of integrin receptor genes, *Itga11* and *Itgb1*, by the midline suture after BAPN treatment.

Defects in collagen fibrillogenesis that we observed with TEM are predicted to decrease the resistance that the collagen network can exert against cellular contractility of adherant mesenchyme and, therefore, soften the tissue’s stiffness. Therefore, we measured the apparent Young’s Modulus of E15.5 and E17.5 BAPN treated embryos and found that, like WT tissue, embryonic midline E15.5 and 17.5 sutures are orders of magnitude softer than flanking frontal bones (compare E15.5 BAPN, Mean = 50.31 Pa ± 2.59 SEM, E17.5, Mean = 226.3 Pa ± 226.3 SEM, and E15.5 bone, Mean = 9077 Pa ± 4094 SEM) Figure 6 E. However, while E15.5 BAPN treated sutures were significantly softer than that recorded in WT, E17.5 BAPN treated reached bulk tissue stiffness comparable to that found in WT at that stage Figure 6 E’. These findings suggest that collagen cross-linking is required for correct suture stiffness, and is a necessary feature of suture morphogenesis. Further, these data provide *in vivo* evidence reinforcing a role for programmed, spatiotemporal control of cellular tension during differentiation and organ formation.

As tissue stiffness is correlated with collagen content in many contexts 11 we asked whether stiffening of BAPN treated sutures towards the end of niche formation could be driven by increased collagen expression in response to reduced LOX-mediated crosslinking. Since Type I collagen is the most abundant collagen in developing midline sutures, we measured *Col1a1* and *Col1a2* expression levels in BAPN embryos relative to WT, using qPCR. Indeed, we find a 15-fold upregulation of *Col1a1* and 4.5 fold increase in *Col1a2* mRNA expression in the midline suture of BAPN embryos Figure 6 D compared to WT E15.5 sutures. As collagen content of the ECM can also influence the production of collagen binding proteins, we asked whether cell adhesion to the altered collagen network was also increased in response to BAPN-induced decrease in suture stiffness. Consistent with mechanosensitive transcriptional feedback of structural proteins predicted by previous reports, we also find increased expression of the most abundant integrin receptors expressed in the midline sutures, *Itga11* and *Itgb1* Figure 6 E. These results suggest that an *in vivo* feedback network can allow developing sutures to compensate for defects in physical structure and force-balance, which, in turn, further suggests that maintaining tissue stiffness is an integral feature of this niche.

Next, we asked whether nuclear shape was perturbed by LOX inhibition by analysing nuclear shapes and neighborhood relationships as before. Therefore, we generated nuclear masked from high-resolution images of BAPN treated E15.5. sutures Figure 7 A-D. While, the distribution of nuclear sphericities were comparable between WT and BAPN treated E15.5 sutures Figure 7 C, topological skeletons of BAPN treated suture cells were significantly shorter and exhibited fewer branches Figure 7 E-G. Combined, these results imply that, while the nuclei of BAPN-treated sutures maintain gross geometric similarity (i.e. sphericity), the nuclei either lose or never acquire the distinctive surface complexity and irregularity observed in early stage (E13.5 and E15.5) WT suture nuclei.

**Figure 7.**
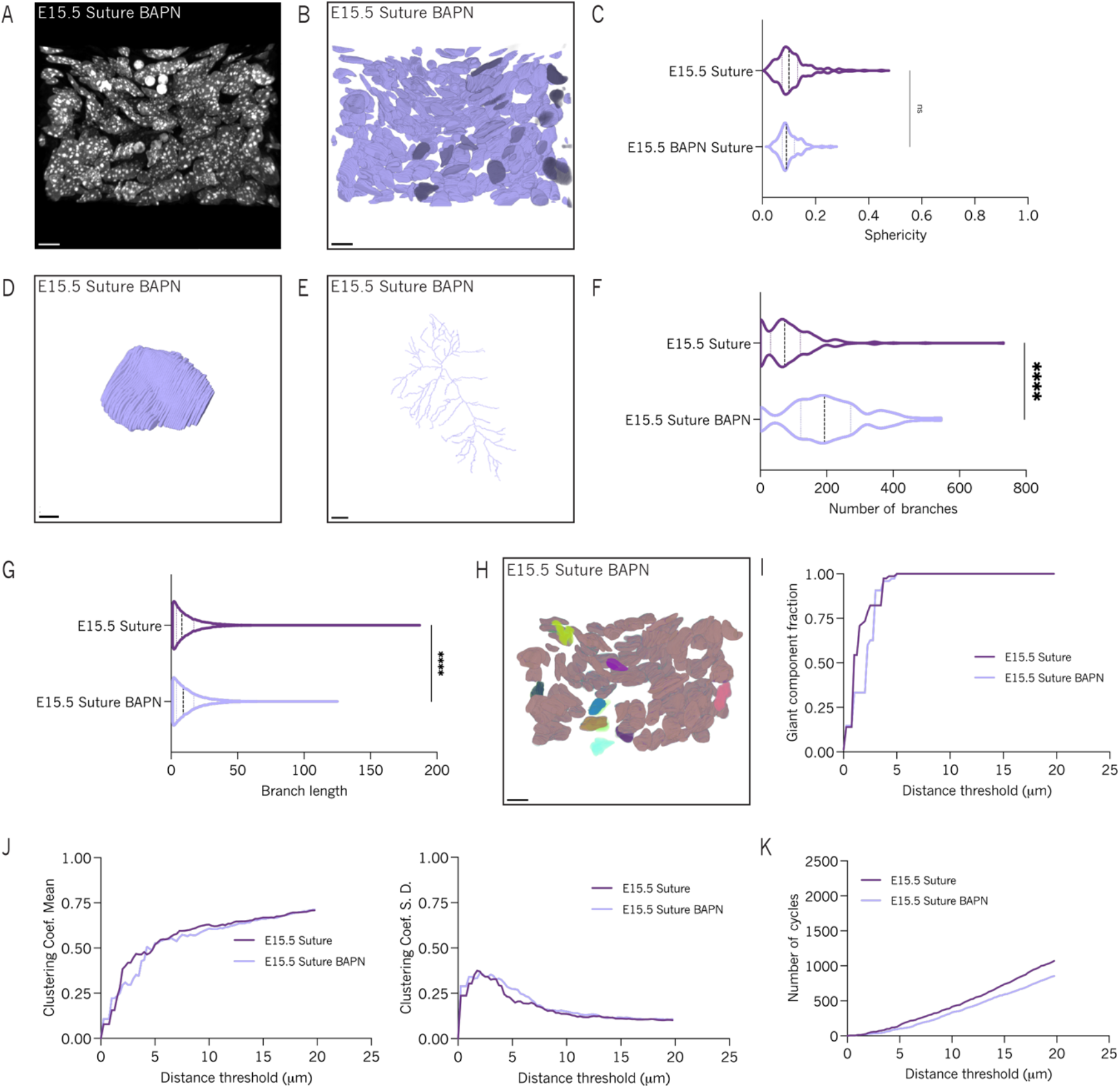
Disruption of collagen fibril formation impacts tissue delineation. A) 3D rendering of confocal z-stack of E15.5 BAPN midline suture. Tissue section was stained with DAPI shown in gray. Scale bar 10 µm. B) 3D rendering of segmented E15.5 BAPN midline suture nuclei. Scale bar 10 µm. C) Violin plot of volume-based sphericity measurements for segmented E15.5 WT (N=2, n=196) and E15.5 BAPN (N=1, n=125) midline suture nuclei. N represents the number of animals, n represents the number of individual nuclei. Mann-Whitney test was applied to estimate p-value. ****p<0.0001. D) 3D rendering of single E15.5 suture BAPN nucleus. Scale bar 2 µm. E) Example of E15.5 Suture BAPN topological skeleton. Scale bar 50 voxels. F) Violin plot of topological skeleton branch number for WT E15.5 and BAPN-treated E15.5 midline suture nuclei. G) Violin plot of topological skeleton branch length for WT E15.5 and BAPN-treated E15.5 midline suture nuclei. ****p<0.0001 H) 3D rendering of nuclear clusters in a BAPN-treated E15.5 midline suture. I) Line plot of giant component fraction for WT and BAPN-treated E15.5 midline suture networks. J) Line plot of mean clustering coefficient for WT and BAPN-treated E15.5 midline suture networks. K) Line plot of clustering coefficient standard deviation for WT and BAPN-treated E15.5 midline suture networks. L) Line plot for cycle basis counts for WT and BAPN-treated E15.5 midline suture networks.

To determine whether BAPN treatment also caused defects in nuclear organisation we generated minimal distance based networks and compared these networks with those found for WT sutures as before. We found that tissue specific cell type clustering was disrupted when compared to WT sutures Figure 7 H and although BAPN-treated networks reach giant component fraction saturation at a point comparable to WT Figure 7 I, the trends observed in clustering coefficient mean and standard deviation were more variable Figure 7 J, K. Further, the cycle count is lower in BAPN-treated sutures than that of WT, which all suggest a change in tissue packing with more isolated, yet evenly spaced cell clusters Figure 7 L. Taken together, the network metrics and lack of spatial separation between presumptive suture, meningeal and dermal cells indicate either a failure to initiate certain steps for suture maturation or a regression from a previously more distinct state.

### Collagen crosslinking is important for enrichment of LMNA/C at the nuclear envelope of suture cells

The transcriptional feedback networks which generate structure in MSC cultures consistently implicate the LINC (linker of nucleoskeleton and cytoskeleton) complex, a specialized nuclear anchoring structure involved in the transcriptional response of the nucleus to mechanical stimuli 41 9 59 60 46 61 47 62 63. While the LINC complex comprises several proteins, including Nesprins (*Syne1, Syne2*), Sun (*Sun1, Sun2*), Lamin (*Lmna, Lmnb1*) proteins mediate LINC mediated mechanoresponse by modulating chromatin dynamics and tissue-specific gene expression 11 39 55 56 57 58 59 38. LMNA also modulates its own transcription where *Lmna* expression and envelope localisation is directly correlated with tissue stiffness, and stiff substrates drive increased *Lmna* expression when MSCs are grown on skeletogenic stiff substrates 8 41 11 46 64 26 9 39 41. Increased matrix stiffness also suppresses LMNA phosphorylation, which is associated with relaxation of the nuclear envelope as well as enhancer binding of phosophorylated LMNA at enhancer sites, including that of *Lmna* itself 64. As we find that developing sutures express LINC complex members and LMNA expression increases over development Figure 8 A, we, therefore, asked whether *Lmna* expression was lower in the relatively soft sutures when compared to bone. Contrary to predictions made by osteogenic MSC experiments 11 41 40, we found that developing sutures had far higher *Lmna* mRNA expression when compared to differentiated reference osteoblasts Figure 8B. Enrichment of *Lmna* expression in sutures was significantly reduced after BAPN treatment indicating that transcription of Lamin A is dependent on the physical structure of surrounding collagen Figure 8 C. As *Lmna* degradation may also be mechanosensitive 9, we investigated if higher *Lmna* expression in midline sutures results in high LMNA localisation at the nuclear envelope, as is seen in osteogenic MSC cultures. Coronal sections of E15.5 heads stained for LMNA/C revealed a gradient of fluorescence intensity at the nuclear envelope which decreased from the midline toward the lateral frontal bones Figure 8 D. To confirm this, we measured fluorescence intensity of Lamin A/C protein in the nuclear envelope of midline suture and frontal bone cells using Fiji. LMNA/C fluorescence intensity values were normalized to the fluorescence intensity of the nuclear stain, DAPI, to confirm increased protein content within suture nuclei compared to bone Figure 8 F. Although, our data suggest an inverse correlation between stiffness and LMNA at the nuclear envelope, we wondered whether LMNA localisation is still determined by ECM organisation as it is in MSC differentiation. Therefore, we also quantified LMNA/C nuclear immunofluorescence in BAPN treated embryos to find a significant reduction in LMNA/C at the nuclear envelope that was significantly different from that of both WT and BAPN treated frontal bones Figure 8 C, D, E. While LMNA concentration within the nucleus is positively correlated with stiffness in other contexts, phosphorylation of that LMNA also impacts sub-nuclear distribution and transcriptional control 9 42 55. Interestingly, when LMNA is phosphorylated it can be found as punctate staining throughout the nucleoplasm, in contrast to the linear enrichment at the nuclear envelope 9 42 55 41 11. In E15.5 midline sutures, LMNA/C is found mostly around the nuclear envelop, whereas in contrast, E15.5 bone exhibited strong LMNA/C puncta across the nuclear interior Figure 8 D’, D’’, respectively. These results suggest that the previously reported correlation between Lamin-A expression with tissue stiffness 11 does not necessarily apply to sutures, if not skeletal differentiation, *in vivo*.

**Figure 8.**
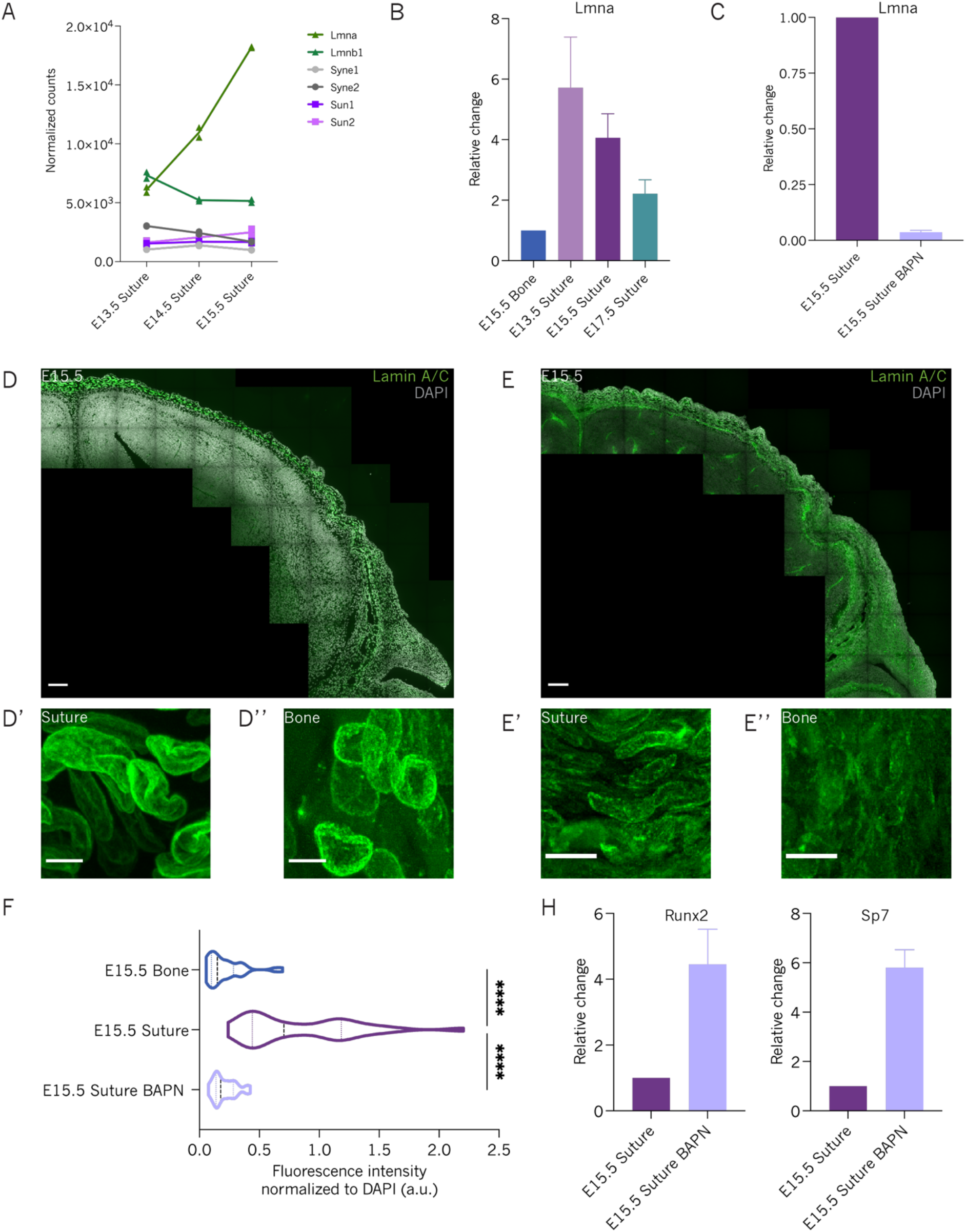
Altered physical properties of the embryonic suture impact nuclear envelop organization and transcription. A) Nuclear envelop genes expressed by the embryonic suture identified by Bulk RNASeq. B) Levels of *Lmna* in the embryonic midline suture (E13.5, E15.5 and E17.5) were measured by qPCR and calculated relative to E15.5 bone. C) Expression levels of *Lmna* in the midline suture are significantly decreased upon BAPN treatment. D) Coronal section of an E15.5 head stained for Lamin A/C (green) and DAPI (gray) shows gradient of Lamin A/C fluorescence intensity along the craniomesenchyme. Scale bar 100 µm. D’) Enlarged view of midline suture region shows smooth distribution of Lamin A/C along the nuclear envelop. Scale bar 10 µm. D’’) Expanded view of the frontal bone shows localization of Lamin A/C around the nuclear envelop as well as foci distributed onver the nucleoplasm. Scale bar 10 µm. E) Coronal section of an E15.5 BAPN head stained for Lamin A/C (green) and DAPI (gray). Scale bar 100 µm. E’) Enlarged view of midline suture region shows reduced distribution of Lamin A/C along the nuclear envelop and foci on thee. Scale bar 10 µm. D’’) Expanded view of the frontal bone shows overall reduced Lamin A/C protein on the nucleus of these cells. Scale bar 10 µm. F) Fluorescence intensity normalized to DAPI was measured for 30 nuclei per conditions. H) Expression levels of *Runx2* and *Sp7* are increased in the embryonic suture after BAPN treatment.

During stiffness induced osteogenesis of MSCs, the balance between LMNA concentration, localisation and phosphorylation is associated with skeletogenic differentiation 41. As loss of collagen crosslinking decreased LMNA content within developing sutures, we asked whether osteogenic gene expression is also altered in BAPN treated sutures. Strikingly, we found a 4 fold increase in the expression of the early master regulator of differentiation, *Runx2* and 6 fold increase in expression of the osteogenic commitment gene, *Sp7* Figure 8 H 82 83. Therefore, interactions with the collagen network control LMNA dynamics within the nucleus that are required to inhibit osteogenesis and maintain SuSC potency, albeit through an inverse correlation to that previously found in MSCs Figure 9 11 64.

**Figure 9.**
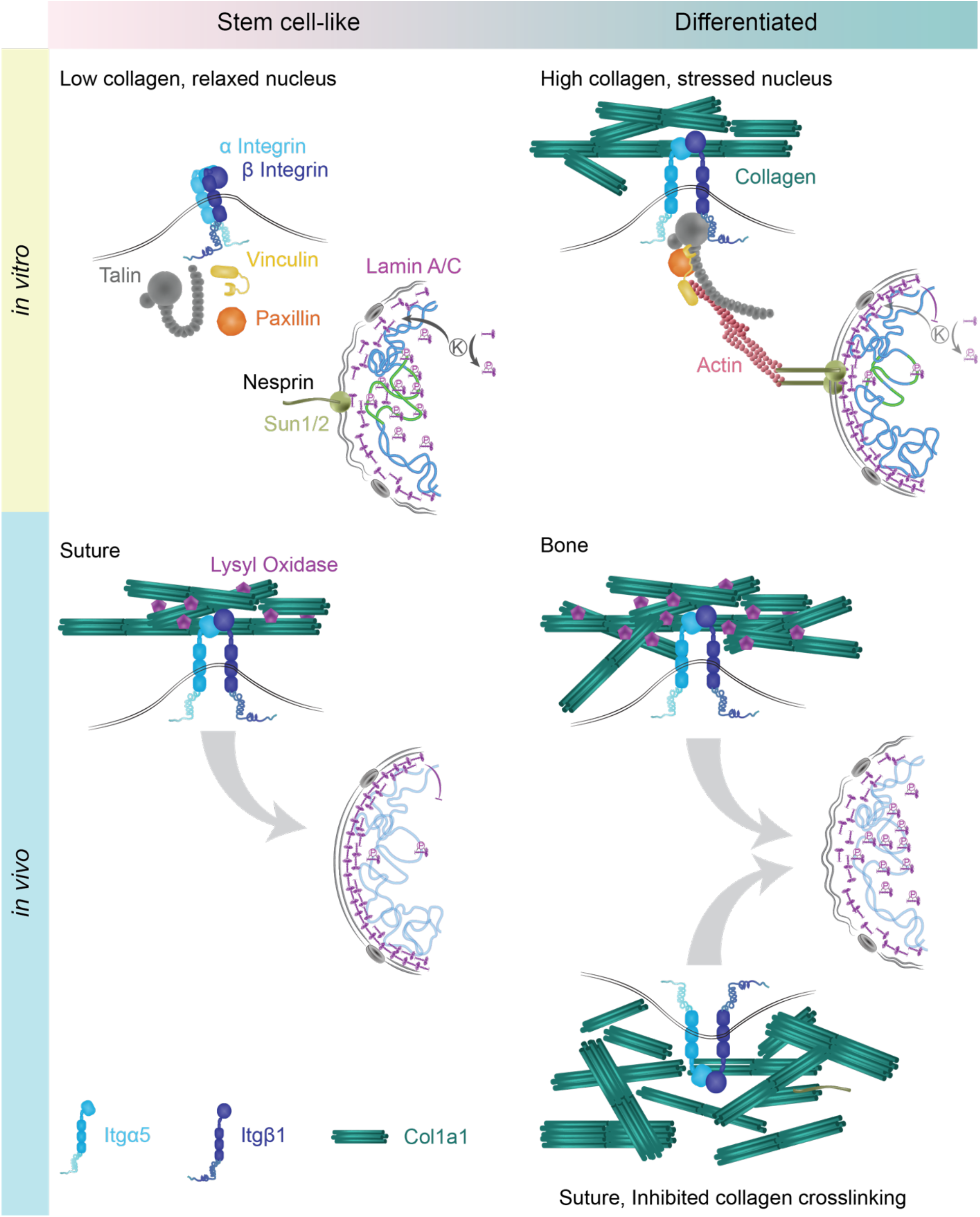
Diagram summarising the nuclear response to extracellular collagen *in vitro* and *in vivo*. In the absence of extracellular collagen, integrin heterodimers remain in a closed confirmation, promoting low nuclear tension and LMNA/C phosphorylation in undifferentiated stem-cell like cultures. In contrast, high extracellular collagen exerts force on the nucleus through integrin mediated actin contractility which increases LMNA at the nuclear envelope. While sutures form a stem cell niche, collagen crosslinking stimulates enrichment of lamin at the nuclear envelope although transcription of osteogenic genes are inhibited. In contrast, LMNA phosphorylation appears increased in response to the increase in crosslinked collagen of developing bone *in vivo*. In sutures where collagen crosslinking is inhibited, LMNA phosphorylation is promoted and osteogenic genes are upregulated.

## Discussion

Craniosynostosis is an example of a relatively common pathology that benefits from *in vitro* cultivation of skeletal stem cells for implantation and, in some cases, also from engineered bones 84 85 86. As an example, Apert’s patients present both synostosis and incomplete bone formation at the midline 87 88 89. Just as *in vitro* osteoblast experiments often begin with the MSC, so too should experiments that aim to understand the structural regulation of skeletal stem cell fate balance *in vivo*. By investigating these sutures as they are established during midgestation, an opportunity arose to unpack how their structure is regulated to allow for stem cells to reside in a particularly complex and stiffening extracellular environment. While craniofacial movement later canalizes suture-specific collagen structure, SuSCs must still remain stem cells as a complex ECM emerges, while also experiencing dynamic forces from the expanding cortex underlying the cranial mesenchyme.

Our data suggests that suture mesenchyme undergoes dramatic transcriptional changes in early development, far earlier than sutures are often considered distinct, at E15.5. At this stage, we find that Collagen 1α1 and 1α2, also characteristic of intramembranous bone, is already complex but must be regulated to maintain cell organization and transcription. This early role for LOX mediated cross-linking was surprising, given that sutures can dynamically realign in response to changes in force balance across the tissue (e.g. Pkd2 mutant mice 90), or YAP-TAZ defects 5. Initially, we were also surprised to find that BAPN treated embryos were able to compensate for the loss in apparent Young’s modulus by the end of suture morphogenesis at E17.5. These data are consistent with the emerging picture of tensional homeostasis during organ development 91 and suggest that SuSCs modulate collagen production to ensure that the niche is softer than bone as it forms but is of a particular stiffness, ∼200Pa. These insights could serve as a reference to design future experiments that marry core priority areas in cell biology, such as matrix complexity and dynamic force balance, as regenerative medicine seeks to design protocols that better recapitulate the environments of SuSCs, bone, or other connective tissues 13 15 92. However, further perturbations of the collagen network *in vivo* will be important to confirm and explore its role in suture biology

Biomechanical studies often focus either on cellular geometry, dynamic forces (e.g. cell loading), or substrate stiffness which are inextricable *in vivo* as the tissues’ 3D geometry will provide cell-cell and cell-ECM interactions that determine both cell geometry and its reception of physical force. Just as *in vitro* assays have demonstrated a clear role for collagen mediated regulation of cell shape and the nuclear lamina through actomyosin contractility and tuning of LINC complex proteins, we find that the mechanical properties of the collagen network influence cellular organisation and nuclear geometry alongside gene expression during *in vivo* skeletal development. Our data support the emerging role for LMNA in regulating skeletal differentiation of mesenchyme, although the opposing LMNA localization patterns suggest that this signaling axis is not always proportional to collagen content as has been found in cultured cells.

This contradiction may simply arise from the need of SuSCs and neighboring osteoblast progenitors to protect their genome under extreme forces provided by mesoscale features such as brain curvature and growth through tension, all of which are also influenced by coordinated development of surrounding bones. While neighboring bones may experience greater tension at the cellular scale from the more heavily cross-linked collagen network present, at the mesoscale, these stiffening bones which flank the suture may create a force-imbalance, such as more stretch across a pliant midline. At the cell scale, however, such LMNA enrichment could play a role in generating complex nuclear shapes as localised LMNA can also promote bending of the nuclear envelope 93. Although new tools will be needed to explore such hypotheses, our data demonstrate the utility of cranial sutures in probing the link between mechanotransduction at the cellular level with the mesoscale geometry of developing connective tissues, such as the skull.

Further experiments are also needed to understand, at the subcellular scale, exactly how skeletal gene expression might be modulated by the LMNA dynamics we find in these sutures. Unravelling the transcriptional regulation that LMNA serves in SuSC’s will likely arise from integration of heterochromatin anchoring and the ability of phosphorylated Lamin to interact with epigenetic modifiers and transcription factors. While mutations in some phosphorylation sites (S222) cause precocious expression of some genes, LMNA can also promote enhancer silencing through DNA modifications 94 55 95 96. In skeletal development, the genetic networks downstream of Runx2 are equally complex as Runx2 is also a transcriptional activator and repressor whose activity is partly dependent on nuclear dynamics, (e.g. cell cycle and envelope tension) in addition to nuclear curvature as MSCs grown on concave surfaces upregulate RUNX2 97 98 99 26 100. Further, LMNA knockdown reduces Runx2 chromatin binding while Runx2 can modulate collagen expression and is upregulated in BAPN treated embryos, potentially emphasizing RUNX2 as an important target of LMNA mediated mechanoresponse in skeletal fate balance 41.

The geometries of the matrix, outer cell membrane and nucleus link mechanisms across the meso-, cell and subcellular scales due to the mechanical couplings that arise through integrin and LINC signaling. Meanwhile, the specific topological characteristics of the nuclear nearest neighbor networks, reflecting the particular packing and clustering present in the suture, must also emerge from the action of mechanisms finding root across the scales. In such systems where the coordination, cooperation, and competition of mechanisms at lower scales give rise to apparently complicated geometric and topological signatures at mesoscales, it becomes increasingly clear that traditional biochemical or genomic characterizations are insufficient on their own to correlate physical characteristics with fate. We posit that these highly complex, volumetric mesoscale tissues must be met with metrics that are more sensitive to and more capable of capturing the intricate arrangements and contortions of different ECM components, the outer cell membrane and the nuclei. And while complicated enumeration of these arrangements is essentially straightforward, it is also not particularly enlightening. What is truly required is a *minimal basis* of shape description and topological characterization. Such a minimal set of simple shape and topology descriptors would, in essence, allow for a decomposition of the complex geometry and topology of the tissue into its key factors which could in turn be associated to underlying molecular mechanism, regulatory pathway, or fate decision (e.g. 101 102). For the suture we have proposed a number of these simplified minimal bases of shape and topology and have demonstrated their effectiveness at accurately characterizing sub-populations within the tissue. At the same time, many of the insights that we report here have been enabled by the manner in which we have been able to decompose the complex three-dimensional spatial arrangements and relative associations.

Ultimately, however, the current situation in many complex mesoscale tissues remains one in which the forest cannot be seen for the trees. And indeed, in the suture, precisely by not focusing solely on molecular cues we gain the distance required to divorce ourselves from the staggering molecular complexity that would otherwise obscure the changes in potency. While the specific shape and topological descriptors we have advanced as providing a partial minimal decomposition of the complex geometry and topology of the suture may fail to play the same role elsewhere, we strongly believe the philosophy of the approach to be eminently and immediately portable to other mesoscale tissues.

Taken together, our data suggest that sutures begin generating a complex extracellular matrix network at a relatively early stage in skull development and that this niche tunes the mechanical properties of its collagen network to regulate gene expression. While we find that this ECM is of similar composition in sutures to that of bone, these niches employ the same collagens to render very different transcriptional outcomes. Further, this reversal is accompanied by an inversion of LMNA enrichment at the nuclear envelope compared to that seen in MSCs on stiff substrates Figure 9. Midline sutures are, therefore, an exciting and potentially powerful model to study the integration of mechanosensitive pathways in the balance between skeletal potency and differentiation *in vivo*.

## Materials and methods

### Animal husbandry

To study the embryonic suture and frontal bone C57BL/6JolaHsd, Osx1-GFP::Cre 83 and R26^mTmG^ 103 mice were used. Genotyping was performed as described in original publications. All procedures were performed in compliance with German animal welfare legislation and in pathogen-free conditions in the biomedical facility of the Max Planck Institute of Molecular Cell Biology and Genetics, Dresden, Germany. Protocols were approved by the Institutional Animal Welfare Officer.

### Lysyl-Oxidase Inhibition

For collagen crosslinking inhibition studies, pregnant Osx1-GFP::Cre females were fed 0.25% BAPN containing food from E11.5, the onset of skull morphogenesis. For embryo collection, pregnant females were euthanized by cervical dislocation and embryos were collected for downstream analysis. Protocol was approved by the Institutional Animal Welfare Officer.

### Bulk RNASeq

Raw reads were checked for their overall quality using FastQC v0.11.2 104 and mapped to the most recent mouse genome reference assembly (GRCm39). Both read mapping and gene counts estimation based on Ensembl release v103 105 were done using STAR v2.5.2b (-outFilterMultimapNmax 1 -outSJfilterCountUniqueMin 8 3 3 3 -quantMode GeneCounts; 106). The obtained count matrix was filtered for genes with more than 10 counts in at least one of the 6 samples and the counts of the remaining 17027 genes served as input for DESeq2 v1.22.1 107. Differentially expressed genes were identified by applying an adjusted p-value cutoff of 0.01 to FDRs obtained using the independent hypothesis weighting method of the IHW package v1.10.1 108. Identification of enriched GO terms was done using the enrichGO function from the ClusterProfiler package v.3.10.1 109 by adjusting obtained p-values with the Benjamini-Hochberg method and a q-value cutoff of 0.01.

### qPCR

To quantify gene expression in developing sutures and bone, embryos were collected in RNase-free PBS and frontal bones and midline sutures were dissected and snap-frozen in liquid nitrogen and kept at -80 °C. Total RNA was extracted using TRIzol Reagent as described by the manufacturer. RNA concentration and purity was determined by Nanodrop. Single-stranded cDNA was synthetized with Random Primer Mix using ProtoScript II First Strand cDNA Synthesis Kit. Primers for genes of interest were designed using PrimerBLAST (see Table 3). qPCR targets were amplified using in triplicate using FastStart Essential DNA Green Master on a LightCycler 96 Instrument. Relative gene expression was calculated by the ΔCt method using Gapdh as the house keeping gene.

**Table 1.** Antibodies and dilutions.

| Antibody | Manufacturer | Cat. Number | Dilution |
| --- | --- | --- | --- |
| Mouse anti-Lamin A/C | Cell Signalling | 8617S | 1:50 |
| Mouse anti-Twist 2 | Abcam | Ab50887 | 1:50 |
| Mouse anti-Runx2 | Santa Cruz Biotechnology | Sc-390715 | 1:50 |
| Rabbit anti-Klf4 | Invitrogen | PA5-27441 | 1:100 |
| Alexa Fluor 488 Goat anti-Mouse IgG | Invitrogen | A-11001 | 1:500 |
| Alexa Fluor 568 Goat anti-Rabbit IgG | Invitrogen | A-11036 | 1:500 |

**Table 2.** Reagents and equipment.

| Reagent | Manufacturer | Cat. Number |
| --- | --- | --- |
| 3-Amino-propionitril -fumarat (BAPN) | Sigma Aldrich | A3134 |
| 7% Tris-Acetate gel | Invitrogen | EA0358BOX |
| 16% Formaldehyde | Pierce | 28908 |
| Agarose low gelling | Thermo Scientific | A9414 |
| BCA Protein kit assay | Pierce Thermo Scientific | 23227 |
| FastStart Essential DNA Green Master | Roche | 06402712001 |
| Gelatin | Sigma Aldrich | G2500 |
| Glutaraldehyde | Electron Microscopy Sciences | 16220 |
| LightCycler 96 Instrument | Roche | 05815916001 |
| LX 112 - Epon 812 Replacement | LAD Research Industries | 21310 |
| Protease and Phosphatase Inhibitor | Pierce Thermo Scientific | A32959 |
| ProtoScript II First Strand cDNA Synthesis Kit | NEB | E6560S |
| Sucrose | AppliChem GmbH | 141621.1211 |
| Tannic acid | Electron Microscopy Sciences | 21710 |
| TRizol Reagent | Ambion | 15596-026 |
| Vectashield | Vector Labs | H1000 |

**Table 3.**
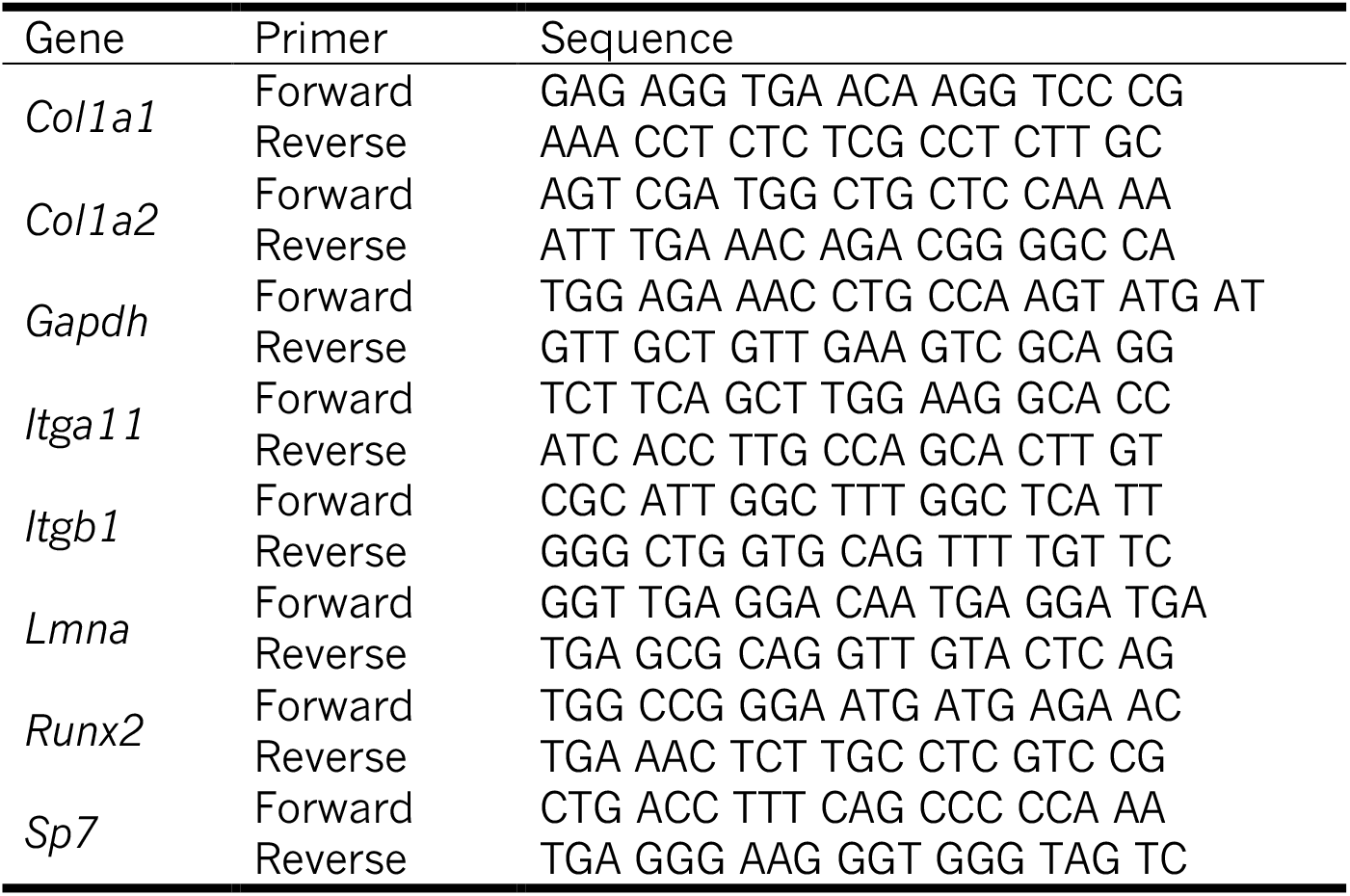
qPCR primer sequences.

### Immunohistochemistry

All immunohistochemistry stainings were performed according to standard protocols. All embryos were collected in cold PBS and fixed in 4% PFA. After fixation embryos were embedded in 15% sucrose/7.5% gelatin and frozen in dry ice. 35 µm coronal sections were collected for nuclear shape and nuclear envelope analysis. Cryosections were stained with Lamin A/C, counterstained with DAPI (1:500, Sigma, 32670) and then coverslipped with Vectashield.

For immunohistochemistry antibody staining 20 µm coronal sections were collected. Antigen retrieval was performed in 10 µm sodium citrate (pH 6.0). Sections were blocked with 10% goat serum in PBS for 1 hour at room temperature and incubated with primary antibodies overnight at room temperature. A list of the used primary antibodies can be found on table1}. Coronal sections were then incubated with secondary antibodies for 2 hours at room temperature, counterstained with DAPI and then coverslipped with Vectashield (Vector Labs, H1000).

### Microscopy

#### Confocal Light Microscopy

For nuclear morphology analysis prepared samples were imaged using Airyscan Super-Resolution Mode acquisition on an LSM-880 (Zeiss). A 63x/1.4 Plan-Apochromat, Oil, DIC (Zeiss) objective was used. Pixel size (0.03 µm), voxel depth (0.2 µm) and laser intensities were kept constant during the acquisition of all datasets.

#### Spinning Disk Microscopy

Samples prepared for antibody staining were imaged using a spinning disc confocal, Andor Revolution WD Borealis Mosaic. A 60x/1.3 U Plan SApo, Silicone (OLYMPUS) objective was used. Pixel size (0.2 µm), voxel depth (0.2 µm) and laser intensities were kept constant during the acquisition of all datasets.

#### Polarized Light Microscopy

For birefringence measurements and collagen orientation analysis 14 µm coronal and sagittal sections were collected, dehydrated in serial steps (25%, 50%, 70% and 100%) of EtOH and coverslipped with Richard-Allan Scientific Mounting Media. Sutures and frontal bones were imaged using an LC-Polscope (on a Ti Eclipse microscope body) with an sCMOS camera (Hamamatsu Orca Flash 4.0) using a 40x 0.65 NA objective (Nikon, MRP50402). µManager 110 was used for data acquisition.

#### Transmitted Electron Microscopy

WT and BAPN embryos were collected as described above and fixed with 5% glutarladehyde/1% tannic acid in 0.1M PBS pH 7.2. Fixed heads were embedded in 4% low gelling agarose and 200 µm sections were obtained using a vibratome (Leica, VT1200S). Sections were post-fixed with 1% osmium tetroxide in water. Sections were dehydrated in serial steps (30%, 50%, 70%, 80%, 90% and 100%) of Ethanol (EtOH), infiltrated with 1:3 EPON LX112/EtOH, 1:1 EPON LX112/EtOH, 3:1 EPON LX112/EtOH and pure EPON LX112. Sections were embedded on Teflon-coated slides with Aclar spacers. Ultrathin cross sections 70 nm) were obtained using an ultramicrotome (Leica, FC7). Sections were post-stained with uranyl acetate and lead citrate and viewed in a Morgagni (FEI) transmission electron microscope, equipped with a Morada CCD camera (Emsis), at 80 kV.

#### Atomic Force Microscopy

For measurements of tissue bulk stiffness embryos were collected in cold 1x PBS, heads were dissected and embedded in 4% low gelling agarose (Sigma, A4018). 2 mm sections were obtained using a vibratome (Leica, VT1200S) and immobilized using glue in a polystyrene-bottom dish (TPP, 93060). Measurements were performed using a Nanowizard IV (JPK Instruments/Bruker) mounted on top of a Zeiss Axio Zoom (Zeiss, .V16). The cantilevers (arrow T1, NanoWorld), modified with 20 µm diameter polystyrene beads (microparticles GmbH), were calibrated by the thermal noise method using built-in procedures of the SPM software (JPK Instruments). Measurements were performed at room temperature (18 °C - 20 °C). Individual force-distance curves were acquired with defined approach and retract velocity (7.5 µm/sec) and with a contact forces ranging from 2.5 nN – 10 nN in order to reach approximately constant indentation depths of 2 µm. At least five specimens were probed for each tissue. The apparent Young’s modulus *E* was extracted from approach force–distance curves by fitting to the Hertz/Sneddon 111 112 model for a spherical indenter using JPK data processing software.

### Image Processing and Analysis

#### Polarized Light Microscopy

The angle frequency distribution of unprocessed PLM orientation images was obtained and used to split each image into several images containing only angles from a specified bin. A morphological operator was applied to the binned images to remove noise from small objects (binary opening process with a 3-pixel wide square kernel). Binned and denoised images were then summed into a single imaged and binarized to create an indexing mask. Original orientation angle measurements were then retrieved using the mask. User-defined midline suture ROIs were then applied to extract the angle distribution.

#### Transmission Electron Microscopy

For collagen fiber size estimation only fibers aligned in the AP axis were considered. Firstly, the outline of at least 380 fibers was manually drawn using *Freehand selection* tool in Fiji 113. Area, aspect ratio and circularity were calculated using *Measure* plugin in Fiji.

#### Confocal Light Microscopy

Lamin A/C fluorescence measurements were done in Fiji. The outline of at least 30 nuclear envelops was drawn using *Freehand selection* tool. Area, integrated density and mean gray value were determined using *Measure*. The same operation was repeated for a background region in the image. Corrected fluorescence (CF) was calculated in the following manner CF = Integrated density - (Area of selected cell x Mean fluorescence of background area). CF of DAPI signal was calculated in the same manner. Lamin A/C CF values were then normalized to DAPI CF.

#### Nuclear Segmentation

Instance segmentation of suture and frontal bone nuclei is a challenging task owing to large variations in nuclei shapes, sizes, density and non-uniform illumination across different spatial regions. Recently, Deep Learning (DL) 114 115 based segmentation methods have achieved remarkable success for microscopy image segmentation. However, such methods need copious amounts of pixel-wise annotated ground truth data for training which is often scarcely available in practice. Especially for 3D datasets, manual annotation is very cumbersome and time consuming since each nuclei has to be annotated across entire volume. To circumvent this bottleneck, we used a 2D annotation and training strategy. We used the open source software called Labkit in Fiji 113 to annotate 417 randomly selected 2D slices of size 512×512 from different 3D volumes (these volumes were not used as part of analysis presented in this paper). We trained a U-Net 116 117 using these annotations and the trained network was used to segment each 2D slice of the 3D volumes used in this paper. The binary segmentation masks obtained from the U-Net are converted to 3D instance segmentations using connected component analysis and any segmentation errors were curated with Labkit.

### Nuclear Sphericity

Due to the complex shapes of suture nuclei, we adapted the classical definition of sphericity 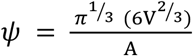 for one that does not depend on a surface area estimation method (i.e. convex hull algorithms or marching cube mesh generation. The resultant definition is 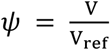, where r_ref_ is defined as 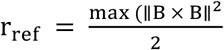 such that B is the boundary set of points for the surface of a given nucleus. This removes the potential under or over estimation of surface area while maintaining the dimensionless range of (0,1) over which sphericity values may vary where a sphericity of 1 indicates a perfect sphere.

### Topological Skeleton Analysis

3D topological skeletons were extracted from all nuclei within the imaging window of each segmented, labeled and object size filtered dataset using previously described methods 71. Skeleton voxels were then classified as either end, chain or branching points depending on the sum of their *N*_2_6 neighborhood. Branches were identified as the continuous collections of chain points situated between either two branching points or a single branching point and an end point. If no branching points were classified for the skeleton, it consists of a single branch (i.e. voxel curve) bounded by two end points. Each branch is assigned a unique identifier (i.e. a scalar). Branch lengths were individually quantified by summing all pixels with the same identifier. The total branch count for each skeleton was also recorded and assigned to the respective nucleus.

### Network Generation and Analysis

Using the boundary sets of individual nuclei, the minimal distance between any two nuclei was determined by searching their distance matrix ∥ *P* × *Q* ∥. Depending on the distance threshold *d* set, edges were created between nuclei. The giant component fraction is defined as the number of nodes (e.g. nuclei) in the largest cluster (i.e. connected components) divided by the total number of nodes in the network. The cycle basis was determined as previously described such that any cycle (i.e. loop) can be created using a sum of parts from the basis 118. The clustering coefficient was calculated as previously described 119.

### Statistics

Statistical analysis was performed using GraphPad Prism ver.9 (GraphPad Software, La Jolla, San Diego, USA). One-way ANOVA is used when multiple quantities are compared, whereas Student’s t-test was used when two samples are compared. Non-parametric tests were applied when normality was not assumed. Normality was measured using the Shapiro-Wilk statistical tests. Values were accepted as significant if the p-value was lower than 0.05. Probability density functions for a normal continuous variable were fitted to datasets for sphericity (Figure 2), skeleton branch counts and branch length (Figure 3) and PLM orientation angle analysis were calculated using the Scipy Statistics package for Python.

## Acknowledgments

We thank the Light Microscopy Facility and Electron Microscopy Facility at the MPI-CBG, in particular to Dr. Dr. Michaela Wilsch-Braüninger and Dr. Weihua Leng at MPI-CBG for assistance with data collection and optimization of imaging protocols. We thank the Scientific Computing Facility at MPI-CBG and in particular Dr. Lena Hersemann for the support with Bulk RNASeq analysis. We would also like to thank Dr. Patrick McCall and Dr. Jan Brugués for engaging in helpful discussion sessions the helped us better interpret our PLM data. We thank Dr. Anna Taubenberger and Dr. Elke Ulbricht for their support with establishing the AFM protocol and for their helpful discussions. We also thank Dr. Anne Grapin Button, Dr. Christian Dahmann, and Dr. Andre Nadler for critical reading of our manuscript. We thank Johanna Lattner for many hours of clicking.

## Author contributions

JMT, DAA and CDM designed research. DAA, and ALC performed research and analysed data. MP and FJ developed deep learning tools. JMT, DAA, CDM and AQR contributed to writing.

## Supplemental material

**Figure supplement I.**
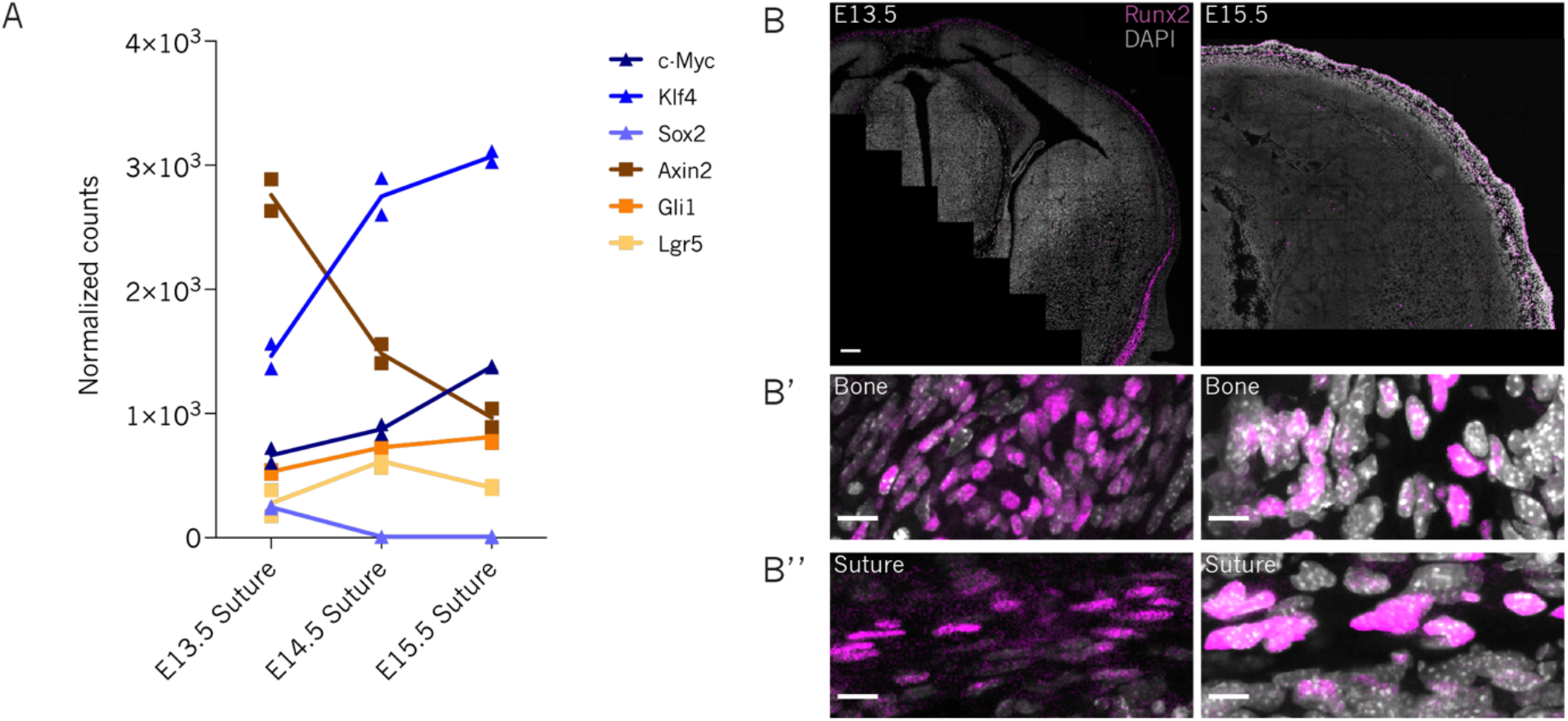
Traditional progenitor markers are broadly expressed in the cranial mesenchyme. A) Expression of stem cell associated markers by the midline suture at E13.5, E14.5 and E15.5. Yamanaka factors expressed by the embryonic suture are denoted by triangle shapes in different shades of blue. Regulators of Wnt or Hedgehog signaling are denoted by squares in different shades of orange. B) Coronal section of E13.5 and E15.5 heads showing Runx2 positive cells in magenta. Scale bar 100 µm. B’) Enlarged view of frontal bone region shows cells positive for RUNX2 at E13.5 and E15.5. Scale bar 10 µm. B’’) Enlarged view of midline suture at E13.5 and E15.5 shows RUNX2 positive cells in the suture. Scale bar 10 µm.

**Figure supplement II.**
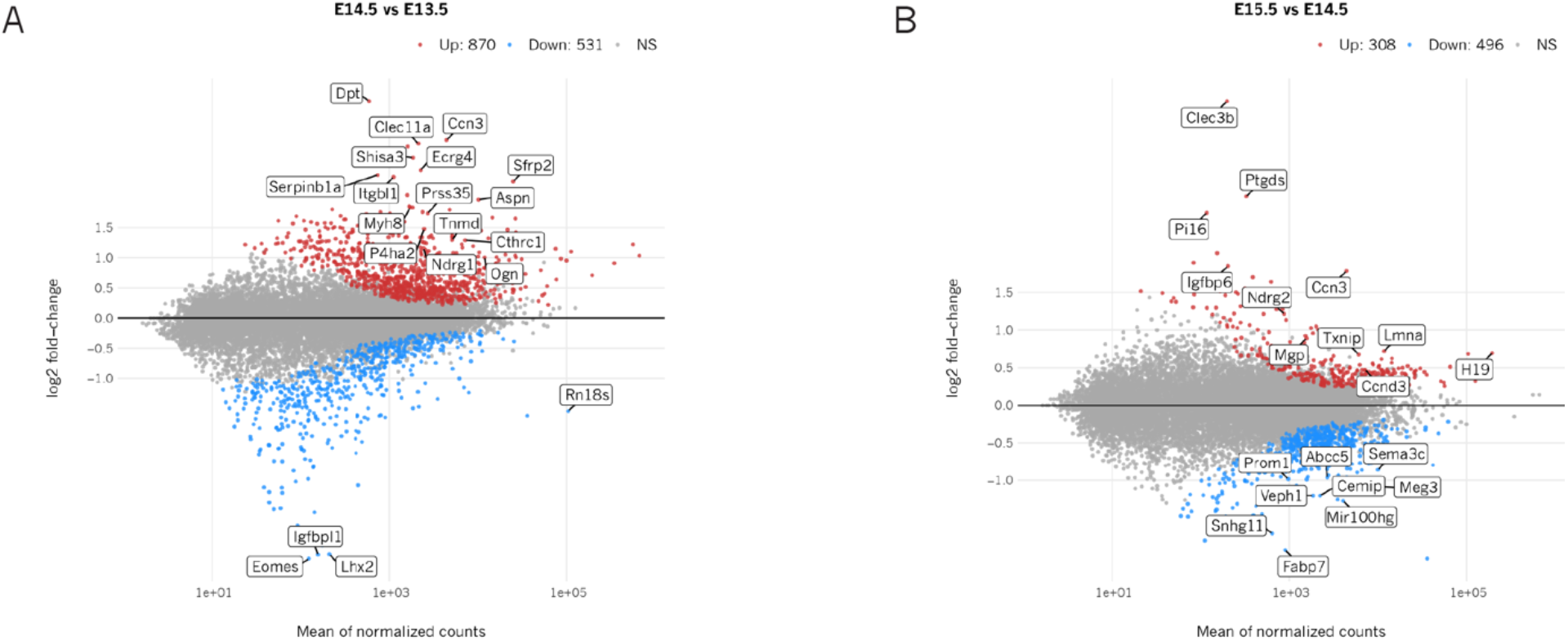
Changes in the midline suture bulk transcriptome during early embryonic development. The log2 fold change for A) E14.5 vs. E13.5 and B) E15.5 vs. E14.5 comparisons is plotted on the y-axis and the average of the counts normalized by log2 is shown on the x-axis. Each gene is represented with a dot. Upregulated genes are shown in red and downregulated genes are shown in blue. Top 20 the of differentially expressed genes per condition have been labeled.

**Figure supplement III.**
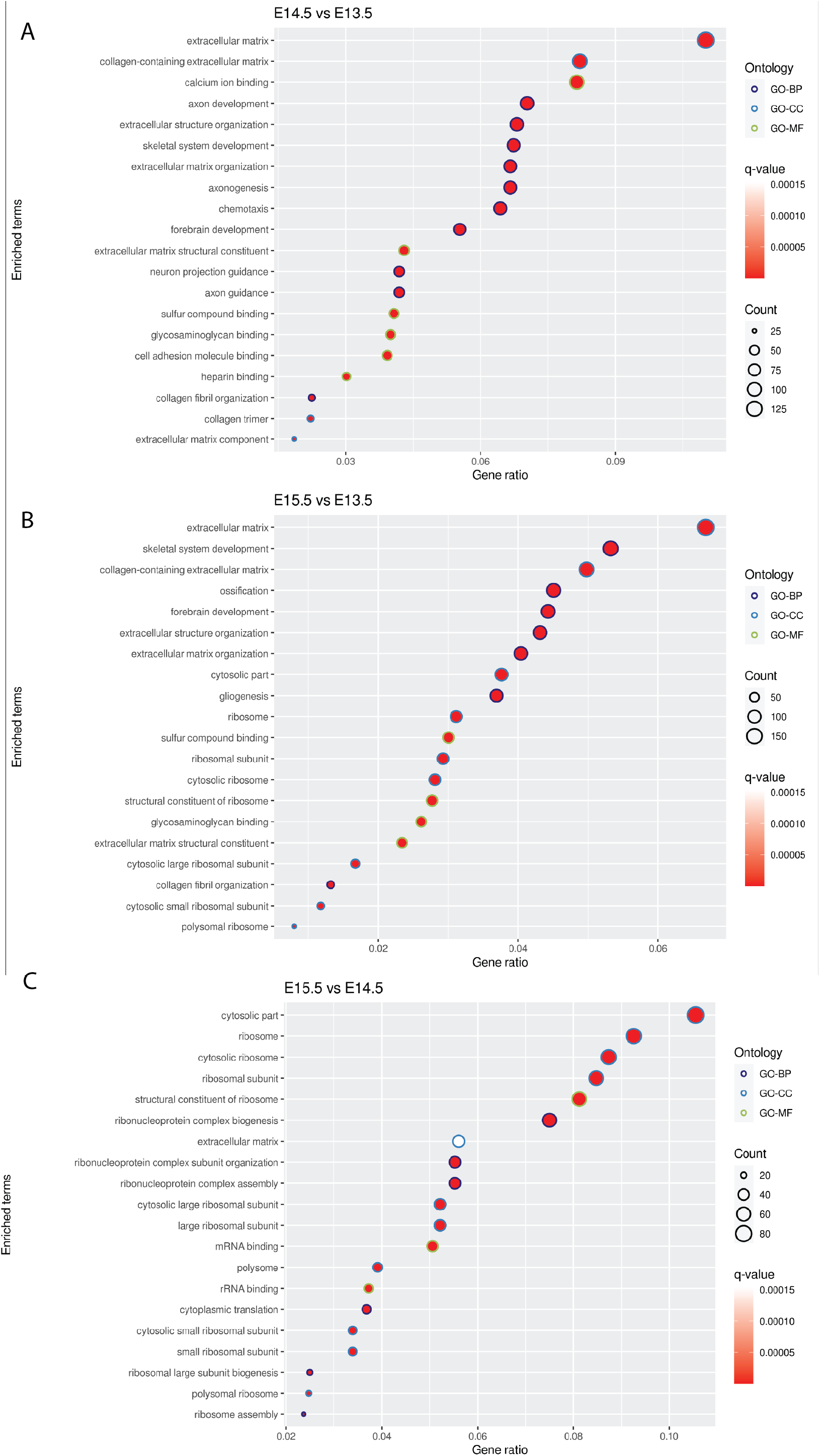
Enrichment analysis. Enrichment terms of A) E14.5 DEGs relative to E13.5, B) E15.5 DEGs relative to E13.5, C) E15.5 DEGs relative to E14.5. GO-BP: Biological Process; GO-CC: Cellular Component; GO-MF: Molecular Function.

**Figure supplement IV.**
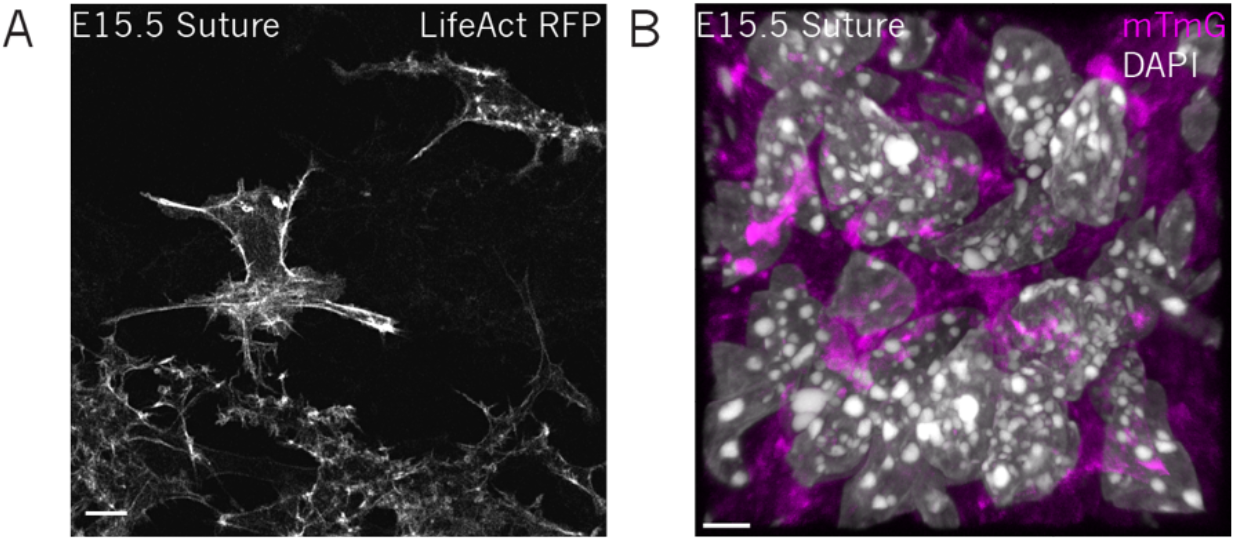
Suture cells have complex membrane morphologies. A) E15.5 suture skullcap transfected with LifeAct-RFP shows that suture cells have long and numerous protrusions and irregular shapes. Scale bar 10 µm. B) Fluorescence imaging of the midline suture from an E15.5 R26^mTmG^ mouse shows diffuse membrane signaling in these cells. Scale bar 4 µm.

**Figure supplement V.**
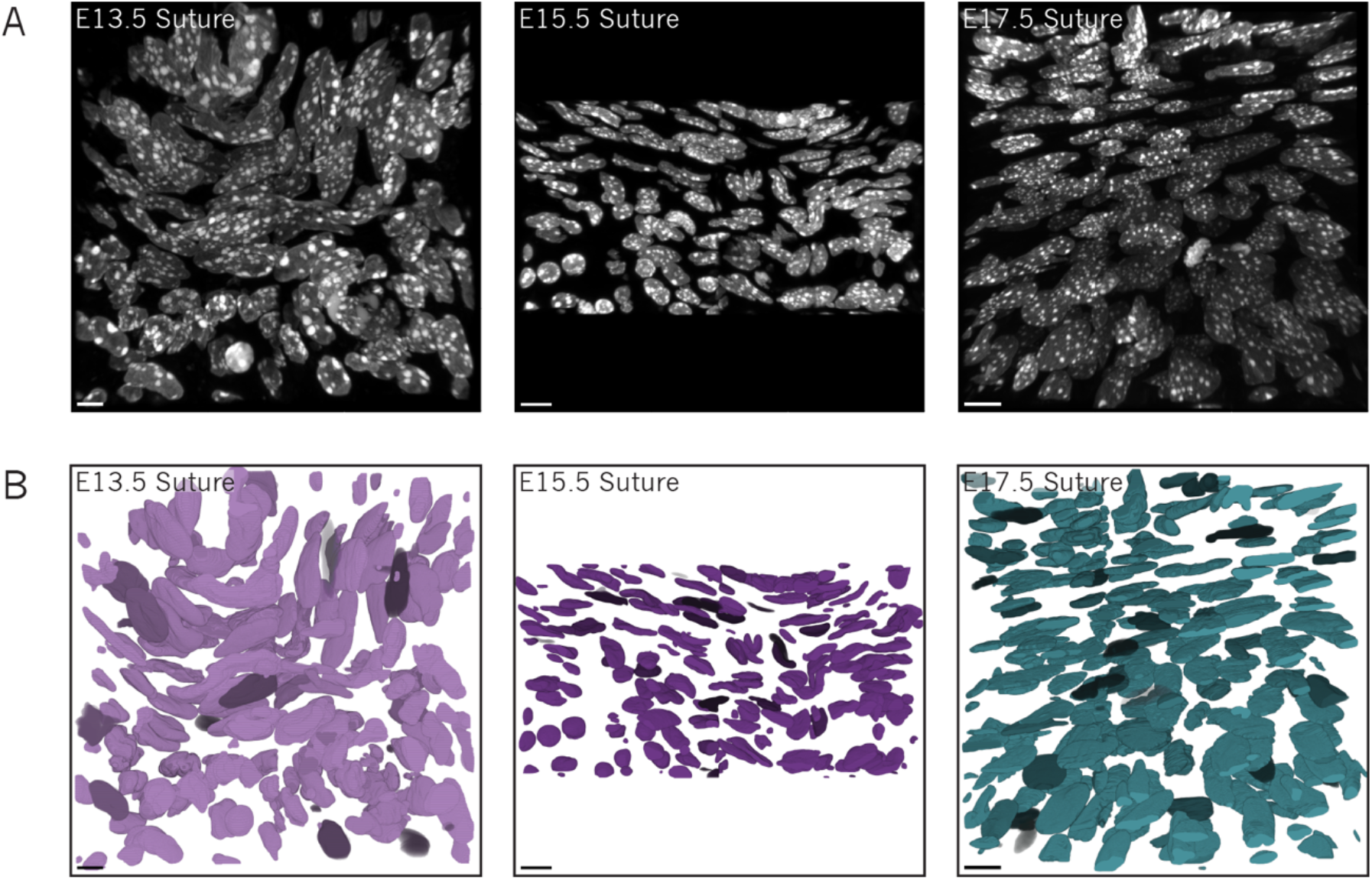
Additional nuclei segmentations. A) 3D rendering of confocal z-stacks of midline sutures nuclei at different developmental stages. Tissue sections were stained with DAPI shown in gray. Scale bar 10 µm. B) 3D renderings of segmented midline suture nuclei at multiple developmental stages. Scale bar 10 µm.

**Figure supplement VI.**
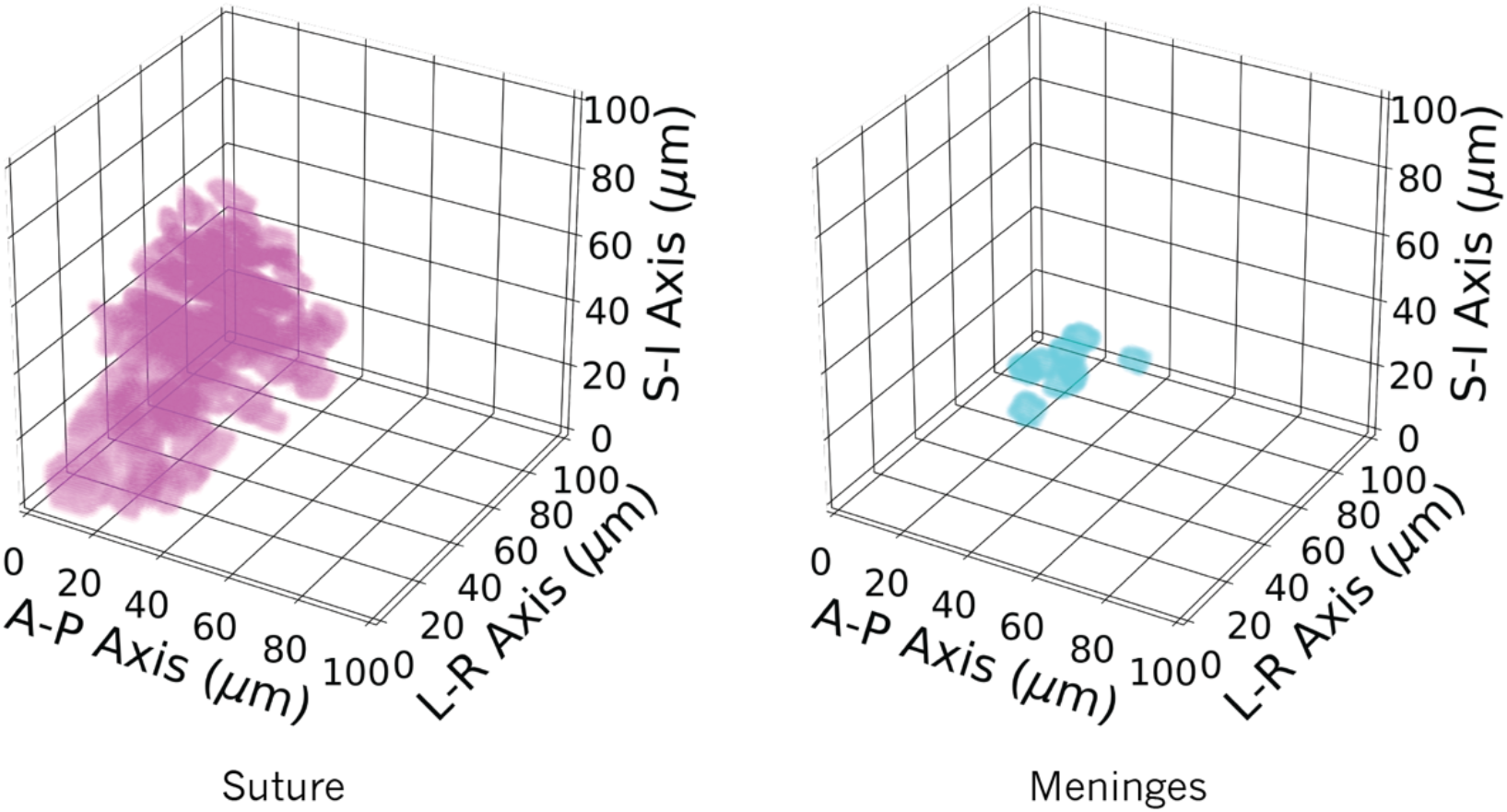
Sphericity thresholding separates cell types. Nuclear sphericity thresholding is sufficient to separate presumptive meningeal nuclei from suture nuclei. Suture nuclei (magenta) and meninges (blue) are identified by their anatomical position.

**Figure supplement VII.**
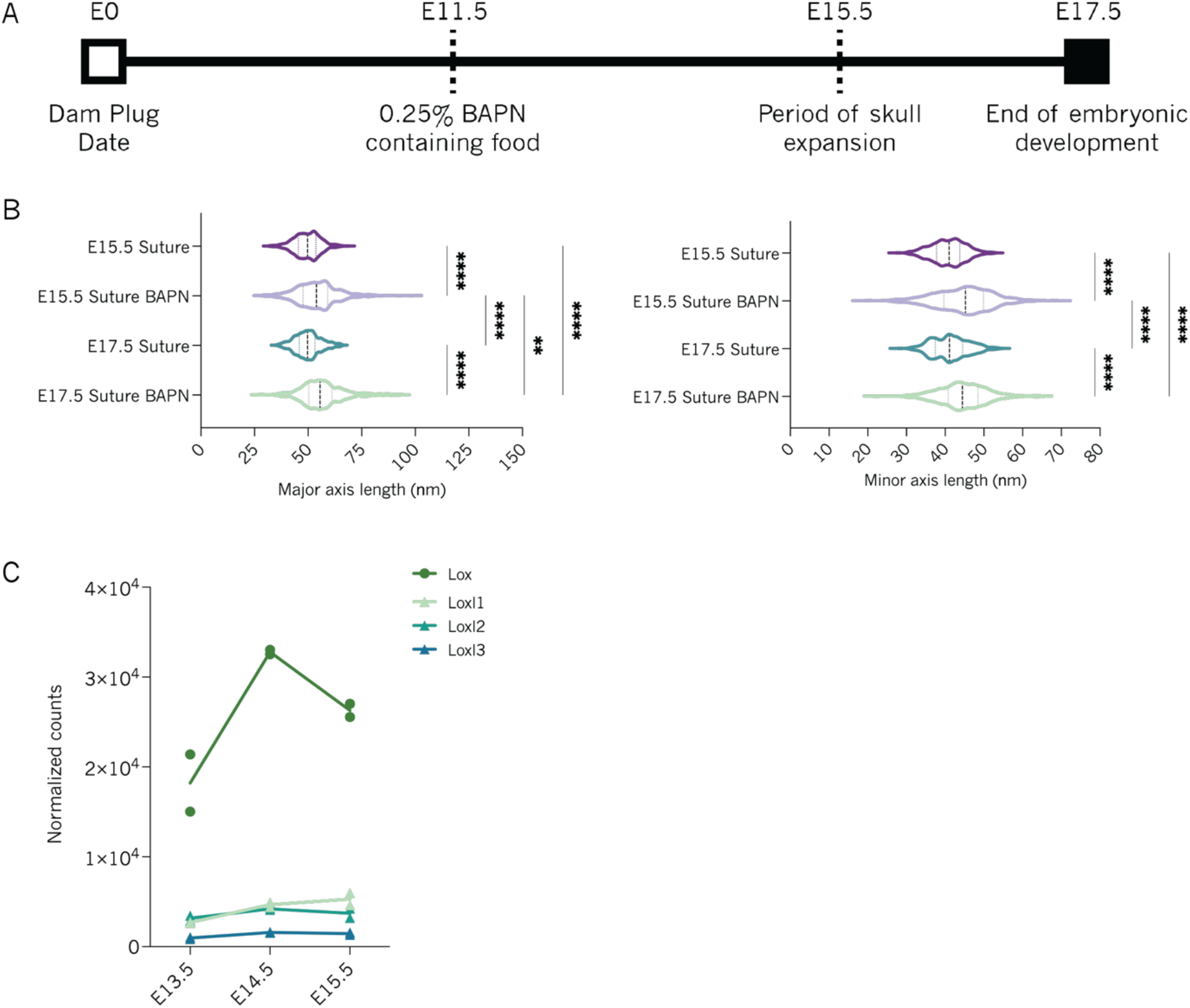
BAPN feeding experiments. A) Scheme for BAPN feeding experiments B) Violin plots comparing major and minor axes length of collagen fibers in E15.5 Suture (N=1, n=401), E15.5 Suture BAPN (N=1, n= 410), E17.5 Suture (N=1, n=269) and E17.5 Suture BAPN (N=1, n=272) embryos. N represents the number of animals, n represents the number of analyzed collagen fibers. Dashed line indicates the median. Kruskal-Wallis test was applied to estimate p-value. *p<0.05, ****p<0.0001. C) Lysyl and lysyl like oxidase genes expressed by the midline suture at E13.5, E14.5 and E15.5.

